# Optimization of a liver Trm cell-inducing mRNA vaccine by reduction of type I interferon response

**DOI:** 10.64898/2026.05.07.723643

**Authors:** Jordan J. Minnell, Mitch Ganley, Ariane R. Lee, Taylah Phabmixay, Kathryn J. Farrand, Olivia K. Burn, John C. Mamum, Emma Lamb, Sarah L. Draper, Susanna T.S. Chan, Olga R. Palmer, Ngarangi C. Mason, Tim Bilbrough, Regan J. Anderson, Benjamin J. Compton, Anton Cozijnsen, Rebecca E. McKenzie, Jochem N.A. Vink, Shirley Le, Dhilshan Jayashinghe, Stephanie Gras, Geoffrey I. McFadden, William R. Heath, Lynette Beattie, Gavin F. Painter, Lauren E. Holz, Ian F. Hermans

## Abstract

CD8+ tissue-resident memory T cells provide rapid frontline protection at pathogen invasion sites, making them attractive targets for vaccine-mediated immunity. We previously developed an NKT cell-adjuvanted mRNA lipoplex vaccine capable of inducing liver Trm cells and sterile protection against malaria in mice. Here, we show that type I interferon (IFN-I) signalling through dendritic cells — not T cells — is a key brake on liver Trm induction by this vaccine. Optimising mRNA manufacturing to reduce immunostimulatory contaminants substantially dampened IFN-I production, boosted antigen expression in the lymphoid tissues, and drove significantly greater Trm accumulation. These enhanced responses translated into superior protection against parasite challenge. Our findings identify DC-intrinsic IFN-I signalling as a tractable target, and mRNA manufacturing quality as a critical and underappreciated lever, for maximising Trm-based vaccine efficacy.

## Introduction

CD8^+^ tissue-resident memory T (Trm) cells are a specialised subset of memory T cells that reside permanently within most tissues and provide local surveillance at pathogen entry sites where they deliver rapid frontline protection against reinfection.^1–3^ Effective generation of Trm cells first requires activation of naïve CD8^+^ T cells upon recognition of cognate antigen, along with a combination of signals that direct effector T cell differentiation and influence their development into either a resident or circulating (effector memory and central memory T cell; Tem and Tcm) memory T cell pathways.^2,4^ These signals include those triggered by T cell receptor (TCR) ligation,^5^ dendritic cell–derived factors (including various cytokines), and inflammatory cues.^6–8^ Following tissue entry, activated CD8^+^ T cells are further influenced by tissue-derived factors^4,9,10^ that drive commitment to a residency pathway, including the shutdown of tissue egress genes like *S1PR1* and *S1PR5* that prevent T cell recirculation.^11,12^

As Trm cells patrol pathogen invasion sites, they are poised to rapidly respond to limit infection. In preclinical models, CD8^+^ Trm cells mediate protective immunity in multiple organs, including the liver,^13–18^ lung,^19–23^ brain,^24–26^ and small intestine,^27^ against a diverse range of viral, bacterial, and parasitic infections. Studies using human tissue samples have identified Trm cells as positive prognostic markers in various diseases, contributing to their control.^28–31^ Given these features, considerable efforts have gone into developing vaccination strategies to elicit effective Trm cell formation as a means of providing long-term protective immunity. mRNA vaccination emerged as a viable strategy for protecting humans from pathogen exposure following the SARS-CoV-2 global pandemic, but localised vaccination using this platform was poor at generating Trm cells at mucosal sites like the lung.^32^

To address these challenges, we developed an mRNA-based vaccine tailored for inducing liver Trm cells, and demonstrated that vaccines targeting *Plasmodium*-encoded antigens provided sterile protection in a mouse model of malaria.^33^ A key determinant of efficacy was the inclusion of a glycolipid agonist, a modified form of α-galactosylceramide (αGC_B_), that recruits help from type I natural killer T (NKT) cells.

Our data indicate that, during the primary immune response, the vaccine is acquired by type 1 dendritic cells (cDC1s), which process and present vaccine-derived components (glycolipid and translated peptide) to activate both type I NKT cells and CD8^+^ T cells. Upon recognition, the glycolipid agonist is presented by DCs to NKT cells that become activated and provide help to CD8^+^ T cells by ‘‘licensing’’ DCs through CD40-CD40L signalling and inflammatory cytokines. Consistent with this mechanism, a prominent feature of the vaccine was the induction of a robust innate immune response, reflecting integration of signals derived from activated NKT cells—together with their downstream cellular cascade—and innate sensing of vaccine components, including the mRNA itself. These signals play a pivotal role in the formation of memory T cells. Specifically, the generation of liver Trm cells was dependent on IL-4, a T cell–modulating cytokine produced by activated NKT cells.^34^ In contrast, type I interferon (IFN-I), likely produced by dendritic cells and regulated by NKT cell activity,^35,36^ impaired the development of liver Trm cells and reduced their protective capacity.^33^

Although IFN-I is essential for the development of robust adaptive immune responses,^37^ its induction during mRNA vaccination can be counterproductive.^38,39^ Excessive IFN-I signalling suppresses cellular protein translation,^38^ thereby reducing antigen production and limiting the availability of peptide–MHC complexes for priming naïve CD8^+^ T cells. In mRNA vaccines, IFN-I responses are primarily triggered by the RNA component itself,^40,41^ underscoring the importance of careful mRNA design. Strategies such as codon and sequence optimization, incorporation of N1-methylpseudouridine [N^1^m(ψ)],^41^ and the use of CleanCap structures^42^ are critical for minimizing potentially harmful innate immune activation while enhancing translational efficiency, stability, and overall vaccine safety.

Here we show that the detrimental impact of mRNA-induced IFN-I is via IFN-I receptor (IFNAR1) expression on dendritic cells (DCs), not the T cells themselves, and that altering vaccine manufacturing to reduce product-like contaminants that drive IFN-I can significantly enhance liver Trm responses to our NKT-cell adjuvanted vaccine. These improved responses were associated with decreased production of inflammatory cytokines and a subsequent increase in vaccine antigen expression in the liver and lymphoid tissues, resulting in more effective T cell responses and increased Trm cell accumulation in the liver. Importantly, vaccines prepared with the improved manufacturing processes provided superior protection against parasite challenge.

## Results

### Liver Trm response to mRNA-LPX vaccine is limited by IFN-I signalling in dendritic cells

We have previously shown that blocking IFN-I signalling with a monoclonal antibody against IFNAR1 at the time of T cell priming with a chicken ovalbumin (OVA)-encoding mRNA vaccine tailored for liver Trm generation resulted in significantly enhanced Trm count.^33^ Here we investigated the impact of IFN-I signalling further in different host settings where IFN-I signalling was abrogated, in this case using a vaccine encoding RPL6 from the rodent malaria-causing parasite, *Plasmodium berghei*. To prepare the vaccine, mRNA was produced by *in vitro* transcription (IVT) with subsequent enzymatic addition of a 5’ Cap-0 and a 3’ polyadenylated (polyA) tail by separate reactions. The mRNA was then complexed with cationic liposomes formed from N-[2,3-(dioleoyloxy)propyl]-N,N,N,-trimethylammonium (DOTMA) and dioleoylphosphatidylethanolamine (DOPE). To promote Trm induction, the NKT cell agonist αGC_B_ was included as an adjuvant in the liposome preparation.^33^ The final liposome/mRNA complexes (LPX) formed vaccine particles of approximately 200 nm in size that were injected intravenously to facilitate interactions with NKT cells enriched in the spleen and liver.

As expected, when compared to an unadjuvanted RPL6 vaccine (RPL6 LPX), a single dose of RPL6 vaccine containing αGC_B_ (RPL6 LPX-αGC_B_) resulted in a significantly higher proportion and number of Trm cells in the liver, defined as activated antigen-specific CD8^+^ T cells (H-K^b^/RPL6_120-127_ tetramer^+^CD44^+^) with a CD69^+^CD62L^-^ phenotype (Figure 1A-C). Increases in central memory T cells (Tcm; CD69^-^CD62L^+^) were also observed, but remained only a minor component of the liver response. Effector memory T cell populations (Tem; CD69^-^CD62L^-^) were not significantly impacted by inclusion of the adjuvant. The response in the spleen was dominated by Tem, which was not impacted by inclusion of aGC_B_, and although splenic Tcm and Trm counts were increased, these populations remained only minor components of the memory response in this tissue. Further characterisation of the dominant liver Trm population compared to liver Tem showed low expression of CX3CR1, and high expression of CXCR6, CD49a, and CD101 (Fig 1D) - markers previously used to describe bona fide liver Trm cells.^13,18,43,44^ Furthermore, tracking of the liver Trm response for 200 days following a prime-boost vaccination regimen with a 4-week interval revealed a half-life of approximately 130 days (Fig 1E), indicating that the vaccine-induced Trm response was long-lived.

**Figure 1:**
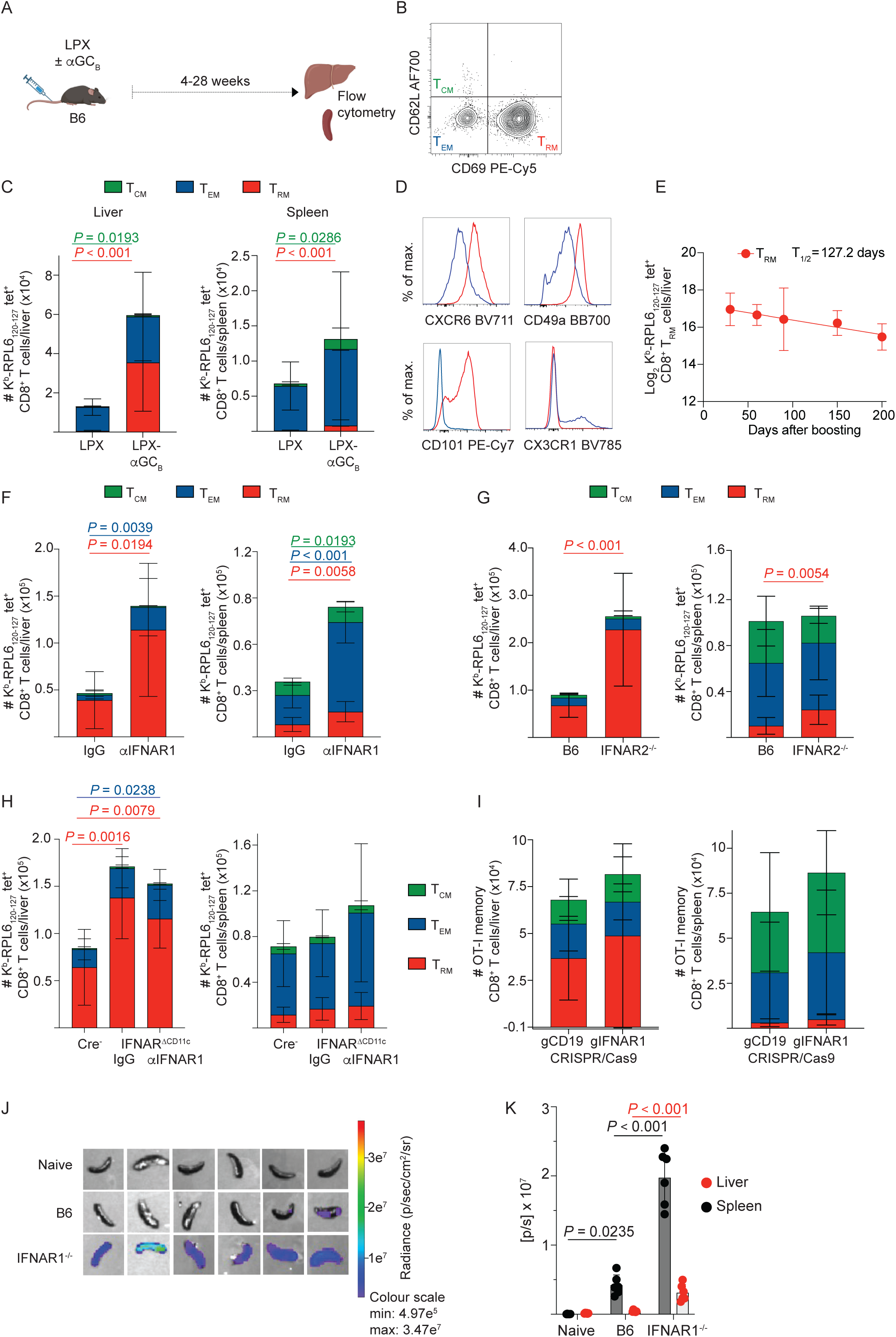
Type I IFNs interfere with antigen expression and liver Trm induction by LPX-. α**GC_B_ vaccines.** (A) Schematic representation of experimental timeline used to assess responses to RPL6-LPX ± αGC*_B_* vaccines. (B) Representative flow cytometry gating strategy used to delineate memory T cell subtypes. (C) Antigen-specific CD8^+^ T cell memory populations in liver and spleen 4 weeks after immunisation, as detected by flow cytometry using K^b^-RPL6_120-127_ tetramers and antibodies for CD26L and CD69 to define memory subtypes: Tcm = CD62L^+^CD69^-^; Tem = CD62L^-^CD69^-^; Trm = CD62L^-^CD69^+^. (D) Representative histograms showing marker expression profiles on antigen-specific Tem (blue) and Trm (red). (E) Liver Trm counts at indicated time-points after two doses of RPL6-LPX-αGC_B_ separated by a 4-week interval (n = 9-10 per timepoint). (F) Memory T cell populations in liver (left panel) or spleen (right panel) 4 weeks after vaccination together with administration of either anti-IFNAR1 (α-IFNAR1) or control antibody (IgG) on day-1 and 1. (G) Memory T cell populations in liver (left panel) or spleen (right panel) of C57BL/6 (B6) and IFNAR2^-/-^ mice 4 weeks after vaccination. (H) Analysis in hosts with Cre-mediated ablation of IFNAR1 expression on CD11c^+^ cells (IFNAR^ΔCD11c^ mice), with α-IFNAR1 or IgG. Cre-negative littermates (Cre-) served as IFNAR replete controls. Data was log transformed and subset cell counts compared by. (I) CRISPR-guides were used to mediate deletion of either IFNAR1 (gIFNAR1) or irrelevant target (gCD19) in OT-I cells. 50,000 donor cells were transferred into B6 mice one day prior to vaccination with OVA-LPX-αGC_B_. (J) Assessment of *in vivo* antigen expression after administration of an αGC_B_ adjuvanted luciferase mRNA LPX. Representative images of isolated spleens 6 h after administration of luciferase-LPX-αGC_B_ in B6 or IFNAR1^-/-^ mice, and naïve controls. (K) Enumeration of luminescent signal in liver and spleen of B6 and IFNAR1^-/-^ mice expressed as mean photon flux per second [f/s] ± SEM. Data are represented as mean□±□S.D.; n = 6. Data in (B) and (D-H) were log-transformed and are represented as mean□±□S.D.; n = 9-10 per group, derived from at least two independent experiments. Memory subsets were compared by student’s t-test. Half-life in (D) was calculated by linear regression. Groups in (I) were compared by one-way ANOVA with Tukey’s post-hoc test. P values are indicated throughout.

To assess whether IFN-I was detrimental to the RPL6-specific liver Trm response, mice were vaccinated either in combination with αIFNAR1 blocking antibodies (transient abrogation of IFN-I signalling) or in a complete knockout setting where mice lacked the IFNAR2 subunit of the type I interferon receptor (IFNAR2^-/-^) (Fig 1F-G). Transient blockade of IFN-I signalling (αIFNAR1) during priming with a single dose of adjuvanted LPX vaccine resulted in enhanced Tem and Trm cell accumulation in both the liver and spleen compared to treatment with an isotype control antibody (Fig 1F). In IFNAR2^-/-^ mice, where IFN-I signalling was abolished, Tcm and Tem cell responses in both the liver and spleen were largely unaffected by the loss of signalling (Fig 1G). In contrast, Trm counts were markedly increased. Together, these findings showed that both transient and complete abrogation of IFN-I signalling enhanced accumulation of Trm cells in the liver.

To determine which cell types were negatively impacted by exposure to IFN-I during establishment of the liver Trm response, responses to the LPX-αGC_B_ vaccine were assessed in mice lacking expression of the IFNAR1 subunit on DCs (IFNAR1^ΔCD11c^ mice). Cre-negative littermates with intact IFN-I signalling served as controls. An additional group of IFNAR1^ΔCD11c^ mice were administered αIFNAR1 during vaccine priming to determine whether IFN-I signalling in other non-DC cell types was involved in generation of memory responses. Vaccine-induced liver Trm responses were significantly enhanced in IFNAR1^ΔCD11c^ mice compared to controls, whereas Tem and Tcm cell responses were unaffected in both liver and spleen (Fig 1H). Unbiased high dimensional analysis of the flow cytometry data confirmed that liver Trm cells were the only T cell population to be enhanced by removal of IFN-I sensing by DCs at priming (Supplementary Fig. 1C). Additional blockade of the IFNAR1 complex in IFNAR1^ΔCD11c^ mice failed to further improve the Trm cell responses compared to isotype treated controls but marginally expanded the liver Tem response (Fig 1H).

These data suggest IFN-I signalling in DCs limits the subsequent liver Trm response to the adjuvanted LPX vaccine. To further address whether signalling in T cells was implicated, vaccination with an OVA-expressing LPX-αGC_B_ vaccine was assessed in mice harbouring transferred transgenic OVA-specific CD8^+^ T cells (OT-I cells) that had *Ifnar1* deleted by CRISPR-Cas9. A control cohort of animals received OT-I cells with deletion of an irrelevant gene, CD19. Memory T cell responses, in particular liver Trm cell responses were equivalent in these mice (Figure 1I). Combined, these data indicate that the detrimental effect of IFN-I signalling on liver Trm cell formation post-vaccination is mediated through IFNAR1 stimulation on DCs, rather than on T cells themselves.

A major role of the IFN-I network is to stall protein translation during viral infection, which limits viral spread between cells. It was therefore possible that the negative influence of IFN-I on liver Trm cell induction was related to vaccine-encoded antigen expression. To assess the effect of IFN-I on mRNA translation and antigen expression, wildtype and IFNAR1^-/-^ mice were injected with luciferase-expressing LPX-αGC_B_ vaccines and assessed with an *in vivo* imaging system (IVIS).

Chemiluminescence, a surrogate of antigen expression, was measured in the spleen and liver 6 h post-vaccination (Fig 1J-K). Significant increases in protein expression were observed in IFNAR1^-/-^ mice compared to wildtype controls, in the liver but more notably the spleen (Fig 1K), suggesting antigen expression is limited by IFN-I induced by the adjuvanted vaccine.

Having identified a detrimental role for IFN-I signalling, we proceeded to investigate whether the induction of IFN-I and other NKT cell-associated cytokines was a result of the NKT cell agonist αGC_B_ or the mRNA-LPX. Activation of NKT cells, with subsequent downstream activation of DCs and NK cells, has been shown to contribute the cytokines IL-4, GM-CSF, IL-12p70 and IFN-γ.^45,46^ The inclusion of αGC_B_ in the vaccine was associated with an increased innate response, as determined by measuring the serum cytokine/chemokine profile over the first 24 h after vaccination with OVA-expressing vaccines (Supplementary Table 1; Supplementary Fig. 2). Significant among the cytokines enhanced with the agonist were CXCL10, CCL2, IL-6, CCL5, TNF, GM-CSF, IFN-γ, IL-12p70, CXCL1 and IL-4.

While IFN-β, which was a very minor component of the IFN-I response to mRNA-LPX, was also increased by the addition of the agonist, IFN-α was unchanged, suggesting that the IFN-I response was driven by the mRNA-LPX itself and that NKT cell activation may enhance this response (via IFN-β induction) further. Considering the major component of the IFN-I response is NKT cell-independent, we sought to modify the mRNA component to minimise IFN-I production and, in turn, enhance liver Trm cell formation post-vaccination.

### Adjuvanted vaccines prepared with optimized mRNA manufacturing processes reduce innate signalling *in vitro*

The inflammatory milieu associated with the adjuvanted vaccine likely represents an integration of signals mediated via activated NKT cells together with innate signals from the vaccine components. This includes the mRNA itself which can stimulate pattern recognition receptors such as the toll-like receptors (TLRs) TLR3 and TLR7, retinoic acid-inducible gene I-like receptor (RIG-I) and melanoma differentiation associated gene 5 (MDA5) as reviewed by Pardi et al.^47^ Given that these innate signalling pathways can culminate in IFN-I release, strategies to limit their stimulation and hence the IFN-I signal may be favourable for liver Trm cell generation. To accomplish this goal, we addressed the synthesis of the mRNA, as this process can induce production of unwanted contaminants that trigger pattern recognition.^41,48–50^

Our “prototype” vaccine used to this point contained mRNA prepared by sequential enzymatic reactions to add the 5’-cap and 3’-poly-A tail moieties to an mRNA transcript generated with native nucleotides.^33^ Not only can this process be costly and time-consuming, but it can also result in batch-to-batch variation in capping efficiency and polyA tail length. As such, several immunostimulants can arise within the transcription reaction, including uncapped RNA, short abortive transcripts and longer dsRNA.^49^ In addition, the inclusion of chemically modified nucleosides such as N^1^m(ψ) into the transcript is a well-defined strategy to reduce innate sensing of IVT-RNA and has been attributed to the clinical success of the COVID-19 vaccines^41,48,50,51^ Therefore, to address whether limiting the IFN-I-inducing capacity of mRNA could enhance liver Trm generation by our vaccine approach, we adapted our mRNA synthesis procedure by: (i) substituting uridine for the modified nucleoside N^1^m(ψ), (ii) directly incorporating a 5’-cap1 structure through an efficient “co-transcriptional” process,^52^ and (iii) encoding the 3’-polyA tail from the DNA template for the IVT reaction to consistently provide a tail of 70 nucleotides (nt). We also assessed the introduction of two post-IVT purification steps; cellulose chromatography to reduce dsRNA contaminants,^53^ and oligo deoxythymidine (oligo-dT) affinity chromatography to enrich for full length polyadenylated transcripts, and remove immunogenic abortive transcripts, using FPLC.

Co-transcriptionally capped and tailed mRNA generated by the adapted IVT method was analysed on an agarose gel and showed a clear band of correct molecular weight that was not remarkably altered by the stepwise introduction of N^1^m(ψ), cellulose chromatography and oligo-dT chromatography (Fig 2A). We assessed the RNA products by ion-pair reversed-phase high-performance liquid chromatography (HPLC) and observed that purification resulted in a more condensed major peak shape, likely reflecting enrichment of full-length transcripts (Fig 2B). We also detected early-eluting material, likely reflecting smaller abortive transcripts,^49^ which were removed by cellulose-or oligo-dT-based chromatography steps, with the latter being superior at removing these unwanted contaminants (Fig 2C). Assessment by ELISA of dsRNA content in the preparations indicated that introduction of modified nucleotides^54^ was sufficient to significantly reduce dsRNA burden (Fig 2D), and these levels could not be substantially reduced further by the chromatography steps.

**Figure 2:**
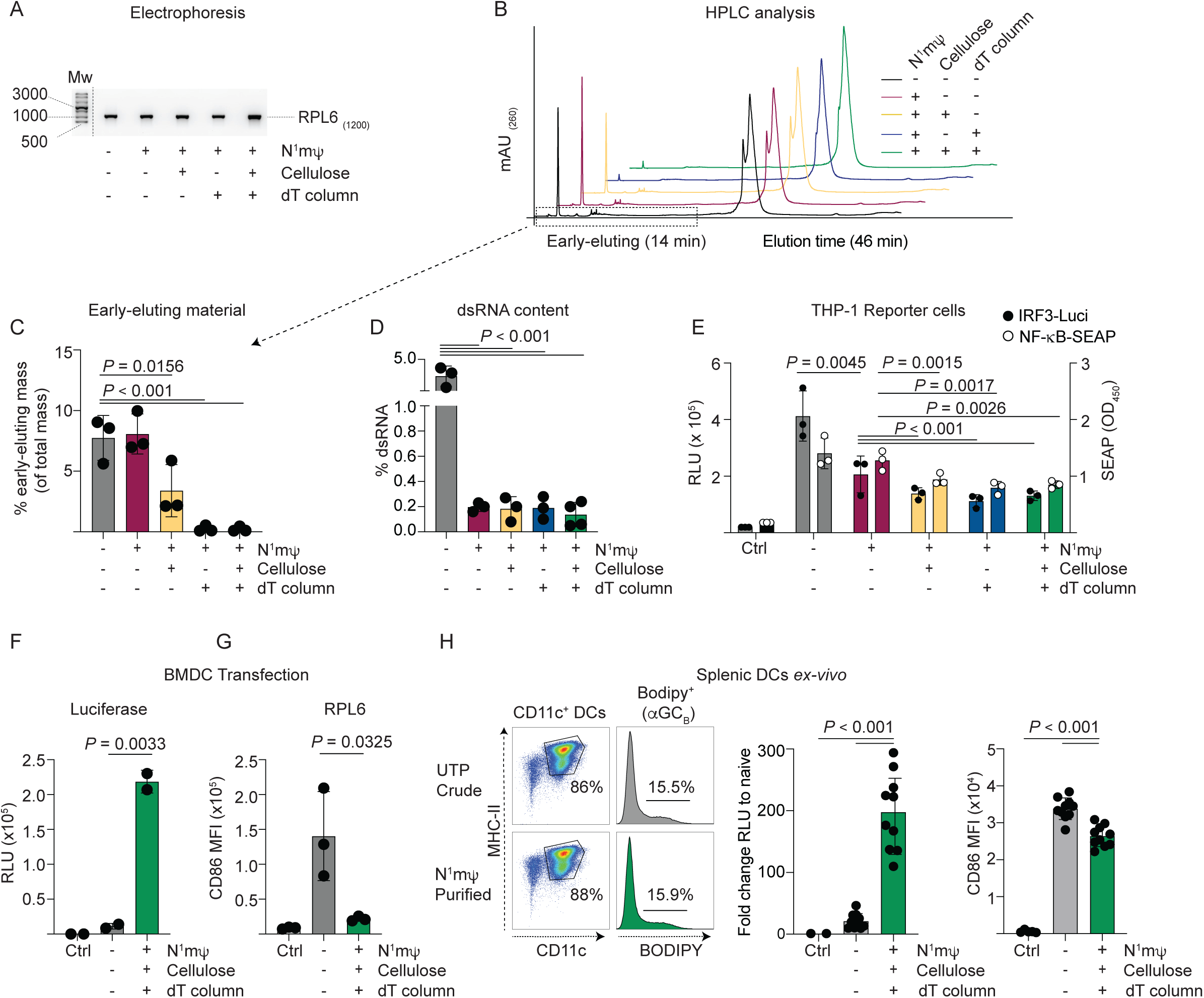
Base-modification and chromatography purification of co-transcriptionally prepared mRNA reduce innate activity of vaccines *in vitro*. (A) Agarose gel of co-transcriptionally capped and tailed RPL6 RNA transcripts following stepwise inclusion of base modification (N^1^m*ψ*), cellulose chromatography, and oligo-dT chromatography as indicated. Lanes contain ∼200 ng mRNA. Image is a composite gel, annotated by dotted line between size marker (NEB ssRNA ladder) and first lane. (B) HPLC analysis of each mRNA preparation depicted by staggered overlays of chromatographs. (C) Area-under-curve analyses of early-eluting products in different mRNA preparations evaluated within gates indicated by dotted box in panel B. Data are represented as mean□±□S.D, n = 3 independent biological replicates (D) dsRNA levels in preparations as measured by ELISA, n = 3 independent replicates (E) Reporter expression from THP1 dual-reporter cell line transfected with LPX vaccines carrying each mRNA cargo, with IRF3 activity expressed as relative light units (RLU) of luciferase reporter expression (black circles, left y axis) and NF-κB activity reported as levels of secreted alkaline phosphatase (white circles, right y axis). n = 3 independent replicates (F) Chemiluminescence signal (RLU) arising from luciferase expression following transfection of BMDCs with luciferase LPX-αGC_B_ vaccines prepared with crude or purified mRNA. Data are represented as mean□±□S.D, n = 2 independent replicates. (H) CD86 expression following transfection of BMDCs with RLP6 LPX-αGC_B_ vaccines. Data are represented as mean□±□S.D, n = 3 independent replicates. (H) Assessment of luciferase LPX-αGC_B_ uptake, luciferase expression and CD86 expession in splenic CD11c^+^ cells enriched from spleen 6 h after injection of the vaccines. Flow cytometry plots on the left show gating of CD11^+^ MHC II^+^ cells (values are mean percentage of CD45^+^ cells in sample) and indicate percentage of cells that acquired BODIPY, a moiety on the αGC_B_ molecule. Graphs shows fold-change in RLU over background in naïve animals (left) and CD86 expeession (right) in gated cells. Data are represented as mean ±□S.D; n = 10 mice per treatment group. Groups were compared by one-way ANOVA with Tukey’s post-hoc test in (C – E; G), or Student’s t-test comparing transfected groups of (F and H). P values are indicated.

However, it was possible that structural differences in mRNA containing N^1^m(ψ) affected the affinity of antibody-based detection techniques.

To evaluate the impact of the adapted mRNA manufacturing steps on innate responses induced by LPX-αGC_B_ vaccines, the dual-reporter THP1 monocyte cell line was used to simultaneously evaluate activation of IRF3, a master regulator of the IFN-I response, and NF-κB transcription factors, which could be involved in production of the other inflammatory cytokines induced by vaccination. Vaccines prepared with the unmodified (“crude”) mRNA induced significant stimulation via IRF3 and NF-κB (Fig 2E). Signalling via IRF3 was reduced by the incorporation of the base modification step alone, although this had no impact on NF-κB. However, introduction of either the cellulose or oligo-dT chromatography steps significantly reduced both signals, but there was no obvious additive impact for combining these purification processes (Fig 2E).

Luciferase-expressing LPX-αGC_B_ vaccines prepared with either unmodified (crude) or N^1^mψ-modified, purified mRNA were evaluated for their capacity to induce antigen expression in dendritic cells (DCs). *In vitro* studies using bone-marrow-derived DCs (BMDCs) demonstrated that vaccines formulated with crude mRNA produced limited luciferase expression, whereas formulation with N^1^mψ-modified, purified mRNA induced significantly higher expression (Fig. 2F). In similar studies, RPL6 LPX-αGC_B_ vaccines prepared with crude mRNA drove marked upregulation of the co-stimulatory molecule CD86 in BMDCs, while N^1^mψ-modification and purification substantially attenuated this response (Fig. 2G). These *in vitro* cultures lacked NKT cells, so no impact of the agonist on BMDC function was expected in these analyses.

Similar findings were observed in splenic CD11c^+^ DCs isolated 6 hours post-injection of luciferase LPX-αGC_B_ vaccines into wildtype mice, where formulation with N^1^mψ-modified, purified mRNA resulted in greater antigen expression and reduced CD86 upregulation, despite similar levels of vaccine uptake (ie. uptake of the BODIPY^+^αGC_B_) (Fig. 2H). The reduction in CD86 was not as marked *ex vivo*, which may reflect DC licensing by NKT cells in the host. Collectively, these data demonstrate that N^1^mψ-modification and chromatographic purification of mRNA reduces innate immune activation while enhancing antigen expression.

### Changes to mRNA manufacturing processes result in enhanced Trm induction correlating with reduced IFN-I

We next evaluated the impact of changes to mRNA manufacturing *in vivo*. Given that reducing IFN-I signalling had been the primary motive for implementing these changes, we first assayed IFN-I cytokines in the serum. This was assessed 8h after B6 mice had been administered LPX-αGC_B_ prepared with the co-transcriptionally capped and tailed crude product, or with mRNA prepared with stepwise introduction of N^1^m(ψ), cellulose chromatography and oligo-dT chromatography (Fig 3A). This showed a stepwise reduction in serum IFN-α with each change, with the final modified and purified preparation leading to a ∼40-fold total reduction (Fig 4B). Similarly, IFN-β levels were also reduced, with base modification and oligo-dT chromatography having the most impact, leading to an overall 100-fold reduction when incorporating all changes (Fig 3C). This latter effect suggested that the NKT cell agonist’s capacity to increase IFN-β still depended on RNA contaminants for induction.

**Figure 3:**
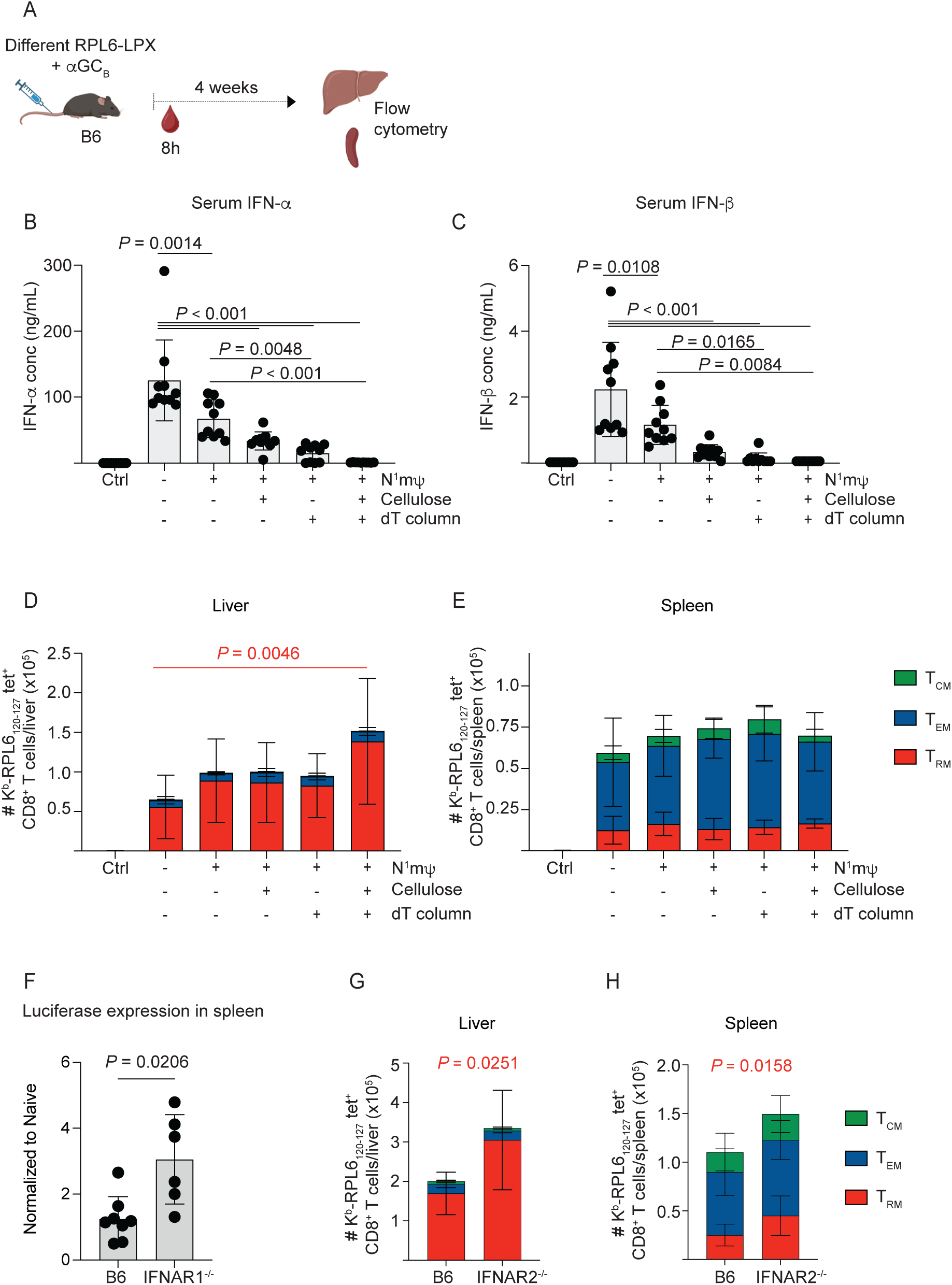
**Base modification and RNA purification enhance liver Trm responses to adjuvanted LPX vaccines**. (A) Schematic depiction of experimental timeline used to assess early (B - C, F) and memory T cell responses (D - E, G - H) to RPL6-LPX-αGC_B_ vaccines (B - C). IFN-I was assayed in serum collected from naive B6 mice and mice vaccinated 8 h earlier with RPL6-LPX-αGC_B_ vaccines outlined in Fig 3. Sera were assayed by LegendPlex assay for IFN-α (B) and IFN-β (C). (D - E) RPL6-specific T cell responses to vaccination were evaluated by flow cytometric analysis of the liver (D) and spleen (E) 4 weeks post-vaccination. (F) Optimized mLuc-LPX-αGC_B_ vaccine was administered to B6 and IFNAR2^-/-^ mice and 6 hr later spleens were removed for imaging by IVIS. (G-H) B6 and IFNAR2^-/-^ mice were vaccinated with the optimized mRPL6-LPX-αGC_B_ vaccine and 28 days later RPL6-specific CD8^+^ T cell enumerated in the liver (G) and spleen (H). Data are represented as mean□±□S.D.; n = 10 in (B – F), n = 8-10 in G and H); derived from two independent experiments. Cell counts were log-transformed (D, E, G, H). Groups were compared by one-way ANOVA with Tukey’s post-hoc test in (B - E), or Student’s t-test in transfected groups of (F - H). P values are indicated.

**Figure 4:**
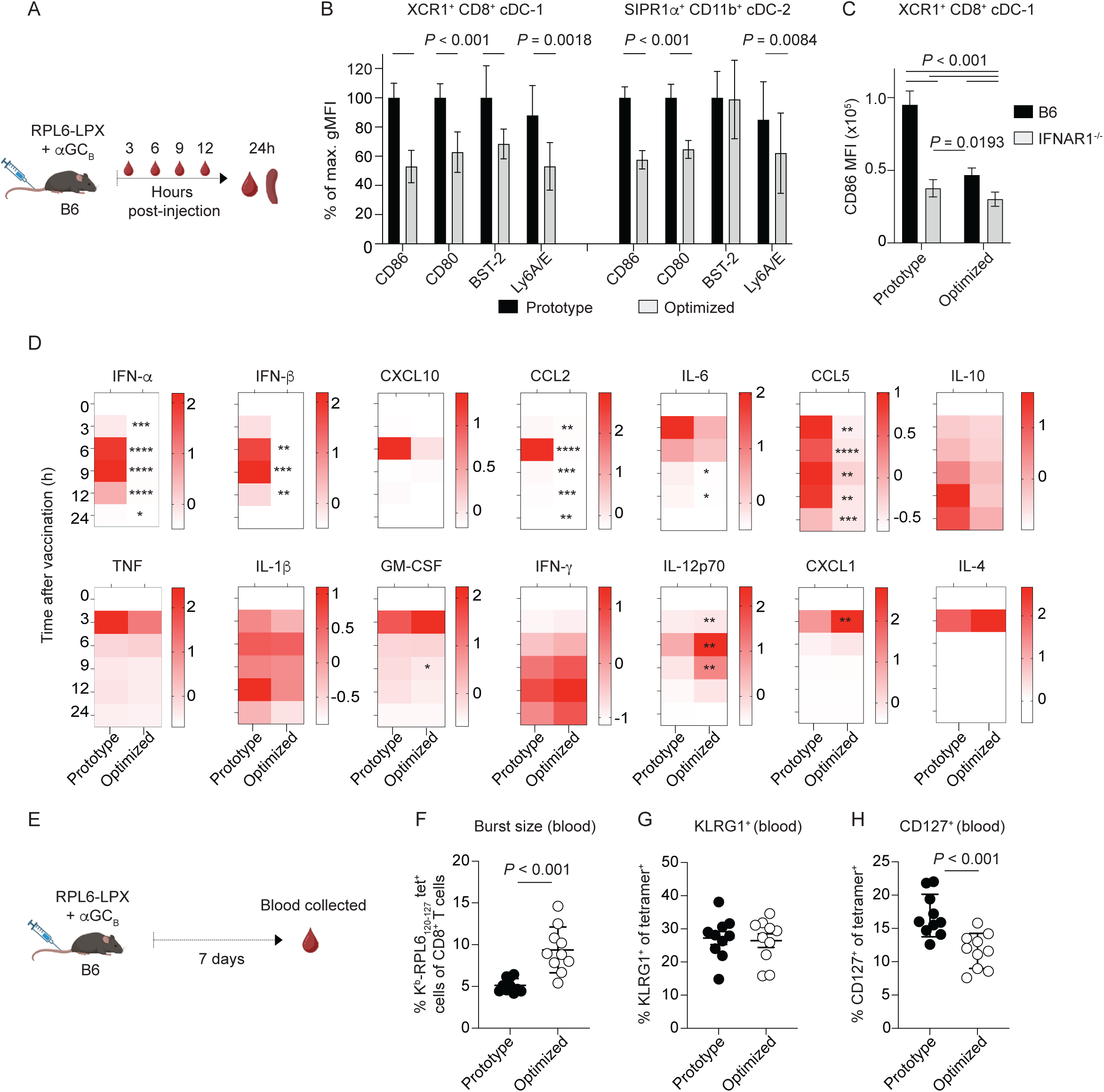
Optimized vaccine alters DC activation profile and reduces inflammatory response during priming. (A) Schematic representation of experimental timeline. Male B6 mice were vaccinated with prototypical or optimized RPL6 LPX vaccines adjuvanted with αGC_B_. Bleeds were assayed for cytokine expression at multiple timepoints within 24 h post-vaccination. (B) At 24□h, splenic cDC1 cells were assessed for expression of CD80, CD86, BST-2 and Ly6A/E expression by flow cytometry. The gating strategy is shown in Supplementary Fig. 6. Data were combined from two experiments and expressed as percentage difference in surface marker expression compared to the average of the prototype vaccine group. (C) Male B6 or IFNAR1^-/-^ mice were injected with prototypical or optimized RPL6-LPX-αGC_B_ vaccines. 24 h later, CD86 MFI was evaluated on splenic cDC1 cells as in panel (B). (D) Serum cytokine/chemokine assay in mice following immunisation with prototypical vaccine with or without αGC_B_. Z-score plots representative of analyte concentration in the serum of mice immunized with optimized or prototypical LPX vaccines at indicated time-points. Lighter colouration represents lower analyte concentrations, and red colouration indicates higher analyte concentration relative to the dataset (analyte concentrations provided in Supplementary Table 2). Data were compared by P values are provided in Supplementary table 2 and also indicated on the Z-score plots: (*P < 0.05, **P < 0.01, ***P < 0.001, ****P < 0.0001). (E) Schematic of experiment used to evaluate peak effector T cell responses in blood. Gating strategy is shown in Supplementary Fig. 5. (F) Percentages of H-2K^b^ RPL6_120-127_ specific cells within CD8^+^ T cells of the blood assessed by flow cytometry on day 7 post vaccination with either prototypical or optimized LPX vaccines adjuvanted with αGC_B_. Proportions of cells expressing KLRG1 (G) or CD127 (H) are shown. Data are represented as mean□±□S.D.; n = 9-10 in (B – D, F - H); derived from two independent experiments. Groups were compared by one-way ANOVA with Tukey’s post-hoc test in (B - C), 2-way mixed effects analysis followed by Sidak’s multiple-comparisons test (D), or Student’s t-test (F - H). P values are indicated.

To assess the impact of the manufacturing changes on liver Trm cell generation, livers and spleens from the vaccinated mice were assessed for memory T cell responses 4 weeks later. We observed a trend for liver responses to increase when the mRNA was less immunostimulatory (Fig 3D). The enhancement in liver Trm response became significant when all the manufacturing steps had been included (Fig 3D), and the IFN-I profile was lowest (Fig 3B-C), resulting in a liver-biased memory T cell response that was >2-fold higher than when the unmodified crude IVT preparation was used (Fig 3D). However, the splenic T cell responses did not differ (Fig 3E).

Given that the mRNA with base modification and purification by cellulose and oligo-dT chromatography led to near undetectable serum levels of IFN-I, we considered whether we would still observe the benefits of this optimized mRNA in IFNAR^-/-^vaccinated mice. To address this point, an αGC_B_ adjuvanted, optimized luciferase-expressing mRNA vaccine was administered to wildtype B6 or IFNAR1^-/-^ mice, with chemiluminescence measured in the spleen 6 h later (Fig 3F). The chemiluminescent signal in the spleen was on average 3-fold higher in the IFNAR1^-/-^mice, indicating that vaccine-encoded antigen expression was still restricted by IFN-I, even from the optimized vaccine. Next, we vaccinated B6 and IFNAR2^-/-^ mice with the equivalent RPL6 mRNA vaccine and assessed the CD8^+^ T cell response in the liver and spleen. Consistent with the average difference in chemiluminescence above, a ∼2-fold increase in Trm cells was observed in the liver and spleen of IFNAR2^-/-^ mice compared to B6 (Fig 3G-H). This was smaller than the effect size previously seen in IFNAR2^-/-^ mice using the prototype vaccine (Fig 1F; ∼3-fold higher Trm count in IFNAR2^-/-^ mice compared to B6). Therefore, although serum IFN-I levels were near undetectable in B6 mice that received vaccines containing optimized mRNA, sufficient IFN-I may have been present in the local milieu at the site of T cell priming to exert a detrimental effect on antigen expression and the liver Trm response.

### Vaccines with optimized mRNA are associated with reduced DC activation, a lower cytokine response, and enhanced T cell burst size

To further examine the impact of mRNA manufacturing processes on early responses to vaccination, we characterised DC activation *in vivo* and systemic cytokine expression following administration of LPX-αGC_B_ vaccines prepared with the prototypic mRNA manufacturing method, or the complete optimized protocol designed to reduce IFN-I (Fig 4A). The phenotype of splenic DCs was assessed 24 h post-injection, focusing on type 1 dendritic cells (cDC1s; defined by expression of XCR1 and CD8) and type 2 dendritic cells (cDC2s; defined by SIRP1α and CD11b).^55^ Both subsets exhibited reduced expression of the costimulatory markers CD80 and CD86 in response to the optimized vaccine compared to the prototype (Fig 4B). Interestingly, this was also associated with reduced expression of the IFN-I regulated protein BST-2 on cDC1 cells, but not cDC2 cells (Fig 5B). Expression of Ly6A/E, a type-I/type-II IFN-regulated protein was reduced on both cDC1 and cDC2 cells. CD86 expression on cDC1 cells was greater in B6 mice that received the prototype compared to IFNAR1^-/-^ mice, demonstrating that CD86 expression is enhanced by the IFN-I pathway (Fig 4C). Accordingly, vaccination with the optimized vaccine led to a smaller difference in CD86 expression between the mouse strains.

**Figure 5:**
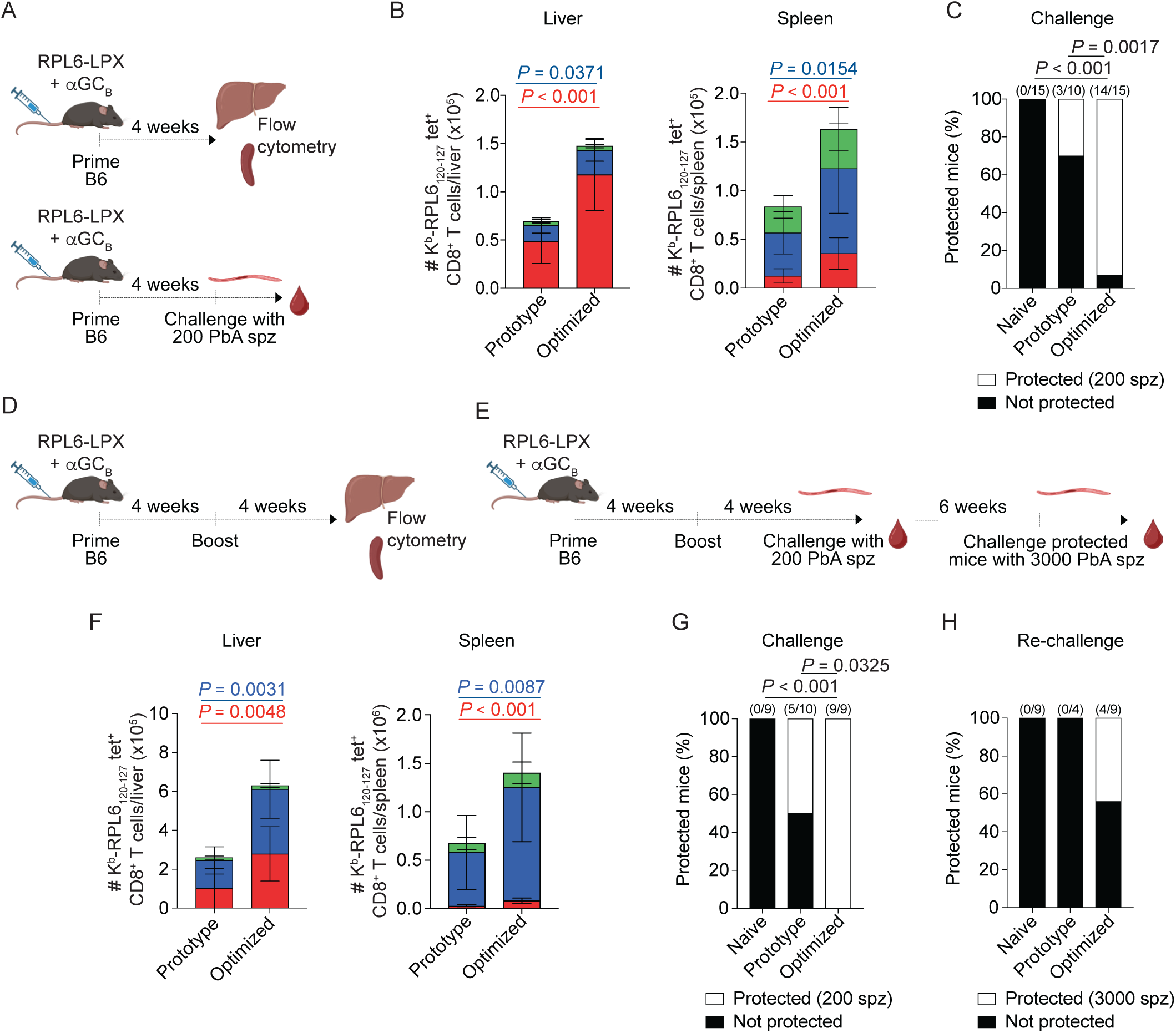
Optimized vaccine improves immunity against malaria. (A) Schematic representation of timeline in (B – C). Mice were administered one dose of prototype or optimized RPL6-LPX vaccine adjuvanted with αGC_B_. Memory responses were measured in the liver and spleen in a subset of mice 4 weeks later. The remaining mice were challenged with a low dose of 200 *P. berghei ANKA* (*PbA*) sporozoites (spz). (B) Enumeration of K^b^-RPL6_120-127_-specific Tcm, Tem and Trm cells in the liver (left) and spleen (right) of mice vaccinated with either the prototypical or optimized vaccine. (C) Percentage of mice protected (white) and not protected (black), as measured by parasitemia in the blood up to day 12 post-infection. Numbers above bars indicate absolute numbers of protected mice to the total number of mice. (D-E) Schematic representation of timeline in (F – H). Mice were administered two doses of either the prototype or optimized RPL6-LPX vaccine adjuvanted with αGC_B_ vaccine with a 4 week interval. (F) Liver (left) and spleen (right) RPL6-specific memory responses 4 weeks post booster vaccination. (G) The remaining mice were challenged with a low dose of 200 spz, and protected mice were re-challenged with a high-dose of 3000 spz. (G-H) Percentage of mice protected (white) and not protected (black) against a 200 spz challenge (G) or 3000 spz re-challenge (H), as measured by parasitemia in the blood up to day 12 post-infection. Numbers above bars indicate number of protected mice to the total number of mice. Data are represented as mean□±□S.D. (B, F); n = 8-15 (B, C, F – H); derived from two to three independent experiments. Data was log-transformed (B, F). Groups were compared by Student’s t-test (B, F), or Fisher’s exact test (C, G, H). P values are indicated.

Vaccine-induced cytokine and chemokine profiles were evaluated in serum over a 24 h period following injection of adjuvanted vaccines prepared with the prototype or optimized mRNA (Fig 4D). As expected, IFN-α and IFN-β levels were substantially lower with vaccines generated with the optimized method, approaching the lower limit of detection. Notably, lower levels of the IFN-I-regulated chemokines/cytokines CCL2, IL-6, CCL5 and GM-CSF were also observed, and substantial reductions in the levels of CXCL10, IL-10, TNF and IL-1β (Fig 4D). The chemokine CXCL1, a chemoattractant for neutrophils, was enhanced. Interestingly, there were corresponding increases in IL-12p70, which is involved in provoking cellular immunity, and (to a lesser degree) IFN-γ, which at this early stage of the response is likely derived from NKT cells and IL-12p70-conditioned NK cells. Expression of IL-4, likely from NKT cells, was unaffected (Fig 4D). Overall, these data point to a general reduction in the pro-inflammatory response, but with enhanced provision of IL-12p70 potentially supporting T cell differentiation, in the context of a significant release of IL-4. Analyte concentration data are provided in Supplementary Table 2.

Enhanced Trm induction using the optimized vaccine was associated with a reduction in provision of co-stimulatory molecules by DCs and a reduced pro-inflammatory cytokine response, but increased IL-12p70 and antigen expression. We therefore assessed whether these conditions could affect early events in T cell priming. To investigate this, CD8^+^ T cell responses induced with the prototypic and optimized vaccines were assessed in the blood at day 7 (Fig 4E). Vaccines prepared with the optimized protocol drove increased expansion of the circulating antigen-specific T cell population, suggesting the stimulatory conditions provided by the vaccine supported overall CD8^+^ T cell priming and enhanced burst size (Fig 4F). IL-12 is known to promote T cell differentiation into short-lived effector cells (KLRG1^+^CD127^-^), rather than memory precursor effector cells (MPECs; KLRG1^-^CD127^+^) which predominantly survive to form memory T cells.^56^ In line with this, although the percentage of antigen-specific T cells expressing KLRG1 was unaltered (Fig 4G), the fraction of these cells expressing CD127, a marker of MPECs, was reduced (Fig 4H). Given that IL-12 may be crucial for memory formation particularly when IFN-I signalling is lacking,^57^ we assessed the T cell response to vaccination in the presence of IL-12p35 blockade. As we previously reported using the OT-I model,^33^ IL-12 is not the key driver of enhanced liver Trm cell accumulation and we confirm this data analysing endogenous T cells (Supplementary Fig. 4). Altogether, this indicates that the greater levels of IL-12 generated by our optimized vaccine is not a driver of the enhanced vaccine-induced Trm cell response.

### Vaccines with optimized mRNA provide superior protection against parasite challenge in a malaria model

In our previous work, we showed that liver Trm cells are the main CD8^+^ T cell subset capable of targeting infected hepatocytes during the liver stage of malaria infection, thereby reducing parasite burden.^13,18,33^ Accordingly, protection provided by the prototypic LPX-αGC_B_ vaccine was abolished when Trm cells were depleted before sporozoite challenge with the mouse pathogen *P. berghei* ANKA.^33^ We therefore assessed whether the improved Trm cell response to the optimized vaccine was able to provide superior protection against challenge in this model. Preliminary studies with the optimized vaccine established that the vaccine dose (5 μg mRNA and 95 μg liposomal lipid content) and adjuvant concentration (80 pmol) used up to this point were both optimal for Trm induction (assessed at 4 weeks after a single dose; Supplementary Fig. 5A-C), and provided sterile protection against challenge with 200 live *P. berghei* ANKA sporozoites (equivalent to about 1 mosquito bite) 4 weeks post-vaccination (animals showing no signs of parasitaemia by day 12 after challenge were regarded as having sterile immunity) (Supplementary Fig. 5D).

In comparing vaccines with this composition, after one dose, the optimized vaccine generated approximately twice as many RPL6-specific CD8^+^ T cells as the prototypical vaccine in both the liver and spleen after 4 weeks (Fig 5A-B). Importantly, a large majority of memory CD8^+^ T cells in the liver were Trm cells – a total ∼2.5-fold increase in counts compared to the response induced by the prototypical vaccine. Vaccinated cohorts were then challenged with 200 live *P. berghei* ANKA. While the prototypical vaccine conferred some sterile protection, as was expected, the optimized vaccine conferred protection in significantly more animals (Fig. 5C). A homologous prime-boost vaccination regimen with a 4-week interval was then assessed (Fig 5D-E). Four weeks after the boost, memory responses were analysed in liver and spleen (Fig 5F). Again, the optimized vaccine generated significantly more RPL6-specific CD8^+^ T cells than the prototypical vaccine in both the liver and spleen. Although the liver Trm component was lower as a proportion of the total memory response for both vaccines when compared to the single dose regimen, the optimized vaccine still induced significantly higher total counts of liver Trm than the prototype. These counts exceeded levels seen in experiments with the single-dose regimen, although a direct comparison was not made. To assess the degree of protection conferred, cohorts of prime-boost vaccinated mice were challenged with 200 PbA sporozoites. Once again, the optimized vaccine provided significantly superior protection, with parasitaemia prevented in all animals (Fig. 5G). To test this protection further, animals protected from the 200 spz challenge were subject to second challenge with the higher dose of 3000 sporozoites (Fig. 5H). None of the mice that received the prototypical vaccine were protected from this high-level challenge, while almost half of the mice that received the optimized vaccine remained malaria-free.

Altogether, the improvements in mRNA manufacturing, with a focus on reducing IFN-I induction, resulted in a vaccine that is efficacious and capable of generating high levels of sterile protection.

## Discussion

In this study, we extend our previous observation that IFN-I negatively regulates liver Trm cell formation by elucidating the underlying mechanism, demonstrating that IFN-I signalling modulates dendritic cells during the T cell priming phase and not the T cells directly. Using our liver Trm cell-inducing adjuvanted mRNA-LPX vaccine,^33^ we demonstrated that in the context of immune signals provided by NKT cells (such as IL-4 and CD40-mediated help),^33,58^ a combination of purification, base modification and co-transcriptional capping/tailing during production of the mRNA component of the vaccine dramatically reduced the systemic IFN-I cytokine response to vaccination and, in turn, enhanced vaccine-induced liver CD8^+^ Trm cell responses.

A number of infection and vaccination studies have shown that IFN-I can support CD8^+^ T cell expansion and survival.^59–61^ However, it has also been shown that IFN-α can suppress CD8^+^ T cell proliferation through activation of the JAK-STAT pathway,^62^ and strong or prolonged IFN-I signalling can impair memory precursor formation.^63,64^ Furthermore, IFN-I is known to inhibit protein translation (as a part of a cell’s anti-viral mechanism),^65^ which could limit antigen availability during T cell priming and constrain clonal expansion.^66^ In previous mRNA vaccine studies, using either lipid nanoparticles (LNPs) or LPX as delivery vehicles, negating IFN-I signalling reduced overall CD8⁺ T cell responses^37,67–69^. In contrast, transient IFNαR blockade during mRNA-LNP vaccination was shown to enhance generation of stem cell–like memory CD8⁺ T cells,^70^ and in a more recent report, transient blockade enhanced cytotoxic capacity and antitumor responses, which was attributed to enhanced antigen load in DCs.^39^ Here, we also observed that transient disruption of IFNAR1 signalling during priming enhanced effector memory T cell populations. This supports the concept that the role of IFN-I signalling can be time-dependent,^71^ with blocking of early IFN-I resulting in improved antigen presentation and subsequent CD8^+^ T cell activation. In terms of the unique capacity for liver Trm formation with our vaccine design, the net impact of IFN-I is inhibitory, mediated solely by signalling receptors on DCs, as the enhanced responses observed in IFNAR^ΔCD11c^ mice were not improved further by αIFNAR blockade, and direct IFN-I sensing by T cells was not required. We also showed that the improvement in Trm response in the absence of IFN-I signalling was correlated with increased antigen expression in DCs. Accordingly, our optimized vaccine, which induced less IFN-I, increased the burst size of antigen-specific CD8^+^ T cells at an effector timepoint (day 7) and the Trm population was preferentially enhanced. This suggests that the unique NKT cell-adjuvanted CD8^+^ T cell priming associated with our vaccine design confers IFN-I-independent memory formation with enhanced Trm accumlation, which is constrained by presence of higher amounts of IFN-I. The enhanced potential for generating liver Trm by our vaccine likely involves Trm precursor formation, with priming predominantly taking place in the spleen,^33^ with these cells having propensity to home to the liver, where local antigen re-encounter with a small amount of LPX-derived antigen can promote Trm formation and persistence.^72–74^ Given that the ablation of IFN-I signalling increased antigen expression in both the spleen and liver, IFN-I may indirectly regulate liver Trm differentiation through controlling antigen exposure at T cell priming and/or during the cumulative exposure in liver tissue. In contrast to our observation that IFN-I can limit liver Trm formation, IFN-I has been suggested to be a key promoter of gut and lung Trm persistence.^75,76^ This may indicate organ-specific dependence on IFN-I, or that additional factors provided as a consequence of NKT cell activation override this requirement with our vaccines.

Previous studies on mRNA-LPX vaccines have shown CD8^+^ T cell responses were dependent on IFN-I signalling,^67,68^ although a profound inhibitory effect of IFN-I signaling was seen upon subcutaneous as opposed to intravenous administration.^67,68^ This again points to the role of IFN-I being time-and context-dependent. A further consideration is the role of CD4^+^ T cells, as IFNAR signalling in CD4^+^ T cells was required for the inhibitory impact on CD8^+^ T cell priming,^68^ possibly impeding the capacity of CD4^+^ T cell to licence DCs. We previously showed the liver CD8^+^ Trm response to be unaffected in MHC II-deficient mice when using our adjuvanted vaccine, with provision of NKT cell-mediated “helper” signals likely rendering conventional CD4^+^ T cell help redundant.^77–79^ Our results here suggest that this NKT cell-mediated help can be qualitatively tuned by reducing mRNA impurities that drive IFN-I, reflected in a general reduction in pro-inflammatory response, but enhanced release of IL-12p70, all in the context of significant release of IL-4 by the NKT cells. While it is known that IL-4 can function as a potent T cell proliferation and differentiation signal,^58^ we have previously shown that IL-4 is essential for the Trm-enriched liver-specific CD8^+^ T cell response to this vaccine.^33^

This study builds on our previous findings that demonstrated that our adjuvanted mRNA vaccine induces liver Trm cell formation that provides sterile protection against malaria infection.^33^ Currently licensed vaccines RTS,S and R21provide partial protection, rather than sterile immunity following malaria infection.^80–83^ Moreover, their efficacy wanes over time, with multiple doses needed for modest protection,^84,85^ presenting significant limitations in the field. With the long half-life of our mRNA vaccine-generated Trm cells and the high level of protection achieved with only 2 doses in a pre-clinical model even after blood stage exposure,^33^ we have shown that mRNA vaccines have great translational potential. The knowledge encompassed in our study detailing the role of IFN-I in liver immunity can be used to guide next generation vaccine designs for malaria and other liver tropic diseases.

## STAR Methods

Vaccines were produced at Ferrier Research Institute and Malaghan Institute of Medical Research. Experimental work (vaccination, tissue processing and flow cytometry) corresponding to Figs. 1A-D, F, H, J-K and Figs. 2 – 4 was also performed at the Malaghan Institute of Medical Research, whereas experimentation (vaccination, tissue processing, flow cytometry, and malaria challenge studies) corresponding to Figs. 1E, G, I and Fig. 5 was performed at the University of Melbourne.

### 1. EXPERIMENTAL MODEL DETAILS

#### Mouse Models

Mice were sex-matched and used between 8 and 12 weeks of age, and were from either the Bioresources Facility at the Department of Microbiology and Immunology, The University of Melbourne, Australia, or the Biomedical Research Unit at the Malaghan Institute of Medical Research, New Zealand. Experiments were carried out under specific pathogen-free conditions, with mice kept at 20–26□°C, 45–65% humidity and a 12 h day–night light cycle. Animals used for the generation of the sporozoites were 4–5-week-old male Swiss Webster mice purchased from Monash Animal Services (Melbourne, Victoria, Australia) and maintained at the School of Botany, The University of Melbourne, Australia. Vaccine studies used C57BL/6J mice (B6) (Jackson Laboratories); OT-I,^86^ IFNAR1^-/-^, (Jackson Laboratories, RRID:IMSR_JAX:028288), and IFNAR2^-/-^ mice (Velocigene).

#### Plasmodium berghei and Mosquito Colony

*Anopheles stephensi* mosquitoes (strain STE2/MRA-128 from BEI Resources, The Malaria Research and Reference Reagent Resource Center) were reared as described previously.^87^ *A. stephensi* mosquitoes (STE2, MRA-128, from BEI Resources) were reared in an insectary approved by the Australian Department of Agriculture, Fisheries and Forestry, which was maintained at 27□°C and 75–80% humidity on a 12 h light–dark cycle. The larvae were bred in plastic food trays (P.O.S.M. Pty Ltd.) containing 300 larvae in filtered drinking water (Frantelle beverages) changed every 3 days and fed with Sera vipan baby fish food (Sera). Upon ecloding, adult mosquitoes were transferred to aluminum cages (BioQuip Products, Inc.) and kept in a secure incubator (Conviron) in the insectary at the same temperature and humidity, maintained on 10% sucrose. The *Plasmodium* species used to raise infectious mosquitoes were *P. berghei* ANKA (PbA) wild-type Cl15cy1 (BEI Resources, NIAID, NIH: MRA-871).

#### Cell Lines

THP-1 Dual cells (InvivoGen) were maintained in RPMI 1640 supplemented with 10% FBS and penicillin/streptomycin (100 U/mL – 100 µg/mL). BMDCs were generated from C57BL/6J mice as described in Method Details below.

#### Ethics Statement

All animal experiments were in accordance with the Prevention of Cruelty to Animals Act 1986, the Prevention of Cruelty to Animals Regulations 2008, the National Health and Medical Research Council (2013) Australian code for the care and use of animals for scientific purposes, or the Animal Welfare Act of New Zealand (1999). The protocols were approved by the Melbourne Health Research Animal Ethics Committee, University of Melbourne (ethics protocols: 25361) or by the Victoria University of Wellington Animal Ethics Committee (AEC14034 and AEC040013).

### 3. METHOD DETAILS

#### mRNA vaccination and antibody blockade

Mice were injected i.v. with 5□µg mRNA in the relevant mRNA LPX vaccine in 200□µL PBS unless otherwise stated. In the instance of IFNAR1 blockade, recipient mice were treated i.p. with anti-IFNAR1 (clone MAR1-5A3) or IgG isotype control (clone MOPC-21) antibodies. Mice were treated with 500 µg on day-1 and 1 mg on day 1. For IL-12p35 blockade, mice were treated i.p. with anti-IL-12p35 (clone C18.2, BioXcell) or IgG isotype control (clone 2A3, BioXcell).

#### DNA template generation

Plasmids for production of enzymatic RNA were amplified linearized as previously described.^33^ For the generation of co-transcriptionally capped/tailed RNA constructs, a G>A modification was made at position +1 relative to the T7 promotor sequence, and DNA templates were generated by PCR.

#### In vitro transcription (IVT) for prototypical vaccine

IVT was conducted from linearized plasmids (see Supplementary Table 3) using the HiScribe T7 High Yield Synthesis kit according to manufacturer’s instructions (NEB). Following IVT, DNA template was degraded by addition of DNase I for 20 min at 37°C. RNA was precipitated in 2 M LiCl and incubated at −20 °C for 20 min, then pelleted by centrifugation at 16,000 × g for 20 min. The pellet was washed with 70% ethanol and centrifuged at 16,000 × g for 5 min. RNA was resuspended in nuclease-free water (NFW; NEB) to a concentration of 1 mg/mL and stored briefly on ice prior to downstream processing.

#### Enzymatic 5’ Capping

IVT RNA was diluted to 0.66 mg/mL in NFW on ice. Capping reactions were performed in 500 µL volumes at 37°C in a thermal block for 1 h and parallelized as required. Each 500 µL reaction contained the following: 50 µL of 10 X capping buffer, 25 µL 2 mM s-adenosylmethionine, 25 µL 10 mM GTP, 0.25 mg RNA, and 25 µL of Vaccinia Capping System (10 U/µL). After 1 h, RNA was precipitated from the reaction in 2 M LiCl at-20°C as above.

#### Enzymatic addition of polyA tail

RNA was diluted in NFW on ice to 0.66 mg/mL. Reactions were conducted in 500 µL volumes at 37°C in a thermal block for 1 h (Eppendorf), parallelized as required. Each reaction contained the following: 50 µL of 10 X capping buffer, 50 µL 10 mM ATP, 0.25 mg RNA, and 25 µL of *E. coli* polyA polymerase (5 U/µL). Following incubation, RNA was precipitated from the reaction in 2 M LiCl at-20°C as above.

#### Co-transcriptional Capping and Tailing

IVT reactions were performed was conducted as previously described^52^ in 200 µL reactions incubated for 2 h at 37°C. DNA template was degraded by DNAse I incubation for 20 minutes at 37°C (NEB) and precipitated in 2 M LiCl. Solutions were frozen at 20°C and pelleted by centrifugation at 23 000 *g* for 20 minutes at 4°C.

#### mRNA Purification

Cellulose chromatography was conducted as previously described.^53^ Chromatography was performed using an FPLC instrument (Cytiva) and Oligo-dT columns from Sartorius. Briefly, mRNA was diluted in 50 mM Tris pH 7.5/0.5 M NaCl and loaded for binding, washed in 50 mM Tris pH 7.5, and then eluted in sterile water. RNA was filtered, adjusted to 0.5 mg/mL, and stored at-80°C.

#### Preparation of Lipoplex (LPX) mRNA vaccines

##### Liposome preparation

Liposomes were produced by an adaptation of the thin-film hydration method.^33^ DOTMA [1,2-di-O-octadecenyl-3-trimethylammonium propane (chloride salt)] and DOPE [1,2-dioleoyl-sn-glycero-3-phosphoethanolamine] (Avanti Lipids) were combined at a 3:1 molar ratio, dissolved in ethanol or chloroform, and transferred to a round-bottomed flask. Adjuvants were added to 0.06 mol% of total lipid (for the standard 80 pmol dose); for dose titration experiments, adjuvant was reduced accordingly. Solvent was removed under vacuum or argon stream and dried further under vacuum. The resulting lipid film was hydrated with UltraPure DNase/RNase-free distilled water (Thermo Fisher Scientific) by overnight incubation at 4 °C to a final lipid concentration of 6 mM. Liposomes were sonicated and extruded through an 11-pass extrusion with a 200 nm polycarbonate membrane using a Mini-Extruder (Avanti Lipids) or 200 nm NanoSizer MINI Liposome Extruder (T&T Scientific Corp.).

##### Lipoplex Complexation and Characterization

mRNA in 100□mM HEPES-buffered solution (pH 7.2–7.5) containing 1.5□M NaCl was complexed with liposomes using a lipid-to-mRNA phosphate molar ratio of 9:1, to a final concentration of 10□mM HEPES, 150□mM NaCl. Lipoplex size was determined by dynamic light scattering using a Malvern Zetasizer Nano ZS (Malvern Panalytical). Lipoplexes were typically 260□nm in size (Z average) with a polydispersity index of 0.2 and zeta potential of 60□mV. Prior to i.v. injection, mRNA LPX vaccine was diluted two-fold in PBS (Thermo Fisher Scientific).

#### Tissue Processing for Flow Cytometry

##### Blood

Blood was either collected from the tail veins of mice in tubes containing 10□U heparin or collected from the submandibular vein punctured with a 4-nm Goldenrod Animal Lancet and collected into 200□μL 10□mM EDTA/PBS. Tubes were centrifuged at 1503 *g* for 4□min, and cells were resuspended in 1□mL RBC lysis solution (QIAGEN). After 20□min at 37□°C, leukocytes were harvested by centrifugation and resuspended in 200□μL FACS buffer (PBS with 5□mM EDTA, 2.5% bovine serum albumin) for antibody staining in a 96-well plate (see Supplementary Table 4).

##### Liver

Livers were collected into 1 ml RPMI containing 2.5% fetal calf serum (FCS) and 10□U heparin. Each liver was mechanically dissociated through a 70-μm cell strainer and washed with RPMI, and the cell suspension was resuspended in 30□ml 35% Percoll (GE Healthcare) before centrifugation at 500 *g* for 20□min at 20–22□°C, with no brake. The cell pellet was incubated in 5□mL RBC lysis solution for 5 min at 20–22□°C before being washed with 35□mL RPMI and resuspended in 1 mL of FACS buffer for flow cytometry. 0.2 mL of the cell suspension was used for antibody staining (see Supplementary Table 4).

##### Spleen - Lymphocytes

Spleen lymphocytes were isolated by mechanically dissociating tissue through a 70-µm strainer, washing in RPMI with 2.5% FCS, incubating cells in RBC lysis solution for 1–2 min at 20–22[°C, washing again and resuspending in 1 mL FACS buffer. 50 µL was used for antibody staining (see Supplementary Table 4).

##### Spleen – Dendritic Cells

For splenic DC analyses and enrichment, spleens were injected at 4–5 points with 0.5□mL IMDM supplemented with 0.3□mg□mL^−1^ Liberase TL and 0.2□mg□mL^−1^ DNase 1, digested for 30 min at 37□°C and then mechanically dissociated through a 70-μm sieve with 20 mL IMDM. Samples were centrifuged at 572 *g* for 4□min, supernatants were discarded, and pellets were resuspended in 2 mL RBC lysis buffer. Samples were centrifuged and resuspended in 1 mL FACS buffer, and 100□μL was transferred to a 96-well plate for staining (see Supplementary Fig. 3).

For DC enrichment, DCs were positively selected by incubation with anti-CD11c magnetic beads according to manufacturer’s instructions (Miltenyi); Purity was confirmed by flow cytometry. For antibody information, see Supplementary Table 4.

##### Antibody staining and flow cytometry

Antibodies used in the study are provided in Supplementary Table 4. Methods for flow cytometry of T cells in liver and spleen samples have been described elsewhere.^88^ For experiments analysing the endogenous T cell response, lymphocytes were stained with K^b^-RPL6_120–127_-specific tetramers for 1□h at 20–22[°C before staining with surface antibodies. Flow cytometry data were collected on an Aurora (Cytek) cytometer using SpectroFlo v.3 software (Cytek) or an LSRFortessa cytometer (BD Biosciences) using FACSDiva v.9 software (BD Biosciences). Flow cytometry data were analysed using FlowJo v.9 and v.10 (Treestar) or OMIQ.

For high-dimensional analysis, spectrally unmixed data were uploaded to the OMIQ platform. Data were scaled and gated to the T cell population of interest and down-sampled to contain 4000 T cells per sample. Dimensionality reduction was performed using uniform manifold projection analysis (UMAP). The optimal cluster number was obtained by elbow plot and clusters were assigned using the FlowSOM algorithm. Differences in cluster abundances between treatment groups were compared by edgeR analysis.

##### THP1 Dual-Reporter Cell Innate Immune Assay

THP-1 Dual cells (InvivoGen) were seeded in 96 well plates at 1×10^5^ cells per well in RPMI, 10% FBS and Pen/Strep at 0.1 mg/mL. mRNA-LPX vaccine samples were added to plate at 250 ng/µL per well, mixed by pipetting and incubated at 37°C, 5% CO_2_ for 24 h. To measure IFN-I activity, 10 µL supernatant was added to 50 µL QUANTI-Luc reagent working stock (InvivoGen) in opaque white 96 well plates and luminescence was read immediately on a VICTOR Nivo Multimode Microplate Reader (Revvity). NF-kB activity was assessed by adding 20 µL supernatant to 180 µL QUANTI-Blue working stock (Invivogen) in clear 96 well plates and incubated for 1 h at 37°C, 5% CO_2_. Absorbance was measured at 650 nm on a VICTOR Nivo Multimode Microplate Reader (Revvity).

##### Bone Marrow-Derived Dendritic Cell (BMDC) Culture and Transfection

Bone marrow was collected from the femur and tibia of C57BL/6 mice into sterile IMDM medium on ice by excision of the bone ends and flushing out with a 25-gauge needle. Marrow was filtered by mechanical dissociation through a 70-µm cell strainer sieve, which was rinsed with sterile IMDM. Cells were pelleted at 250 *g* for 10 mins at 4°C and pellets were lysed with 1 mL of red blood cell lysis buffer (Qiagen) for 2 mins at room temperature and washed once more in IMDM supplemented with 5% FCS. Cells were resuspended to 0.2 million cells/mL and cultured in 6-well plates in the presence of 150 ng/mL of recombinant human Flt3L (BioXCell). Cultures were maintained by changing 40% of the media volume and replenishing the rhFLt3L at days 3 and 6 post-seeding. At day 9, cells were harvested and seeded for transfection at 0.5 million cells/mL in the presence of rhFlt3L. After 24 h (on day 10), 1 µg of LPX vaccine was added to cultures. Cells were harvested by mechanical dissociation 24 h later and stained for flow cytometric evaluation of surface markers (antibody panel described Supplementary Table 4; gating strategy in Supplementary Fig. 6).

##### Assessment of Luciferase Expression (*in vitro and ex vivo)*

Following transfection of BMDCs, or isolation of splenic CD11c^+^ DCs, culture medium was gently aspirated off the wells, and 40 µl of sterile, colourless RPMI 1640 supplemented with 10% FBS and 1 % Pen/Strep (Thermo) added to each well. The Biotium Steady-Luc working solution was prepared in the dark by diluting D-luciferin (10 mg/ml) in assay buffer to give a final concentration of 0.25 mg/mL. 40 µL of working solution was added to each well, and left on a shaker for 5 min at 350 RPM in the dark. Following a further 10-15 min incubation at room temperature in the dark luminescence was measured on a VICTOR Nivo Multimode Microplate Reader (Revvity).

##### High Performance Liquid Chromatography Analysis of mRNA

Sample separation was performed on a DNAPac RP column with 4 μm particles and dimensions of 3 × 100 mm (ThermoFisher Scientific) at a flow rate of 0.35 mL/minute and column temperature of 65 °C. Mobile phase A consisted of dibutylammoniumacetate (10 mM) and triethylammonium acetate (100 mM) and mobile phase B consisted of 50% acetonitrile, dibutylammonium acetate (10 mM), and triethylammonium acetate (100mM). Separation was accomplished by step-gradient with an initial 3 minute hold at 5% B, a 1 minute gradient from 5-25%B, a 15 minute gradient from 25-35%B, a 15 minute gradient from 35-37.5%B, a 12 minute gradient from 37.5-100%B and a 2 minute hold at 100%B. mRNA was detected by UV at 260 nm.

##### Double-stranded RNA (dsRNA) ELISA

dsRNA ELISAs were conducted using an EasyAna Modified dsRNA kit (Vazyme). 100µL of modified A two-fold dilution series of the dsRNA standard (1.5 ng/mL to 0.023 ng/mL) was plated alongside mRNA samples (80 ng, 20 ng, or 10 ng in 100 μL) in duplicate onto the pre-coated kit plate. Following a 1 h incubation at 37°C, plates were washed four times with 300 µl wash buffer, then 100 µL of detection antibody was added and incubated for 1 h at 37 °C. The detection antibody was washed off as before, prior to the addition of 100 µL of enzyme-labelled reagent, then plates were incubated for 1 h at 37 °C. The enzyme-labelled reagent was washed off as before then 100 µL TMB substrate was added and incubated for 15 minutes at 37 °C, followed by the addition of 50 µl 1M H2SO4 to stop the reaction. OD was measured at 450 nm and the level of dsRNA quantified using the standard curve.

##### Multiplex Serum Cytokine Assay

Blood was collected in microvette 500 Z-Gel tubes (Sarstedt) and centrifuged at 4 °C for 5 mins at 10,000 *g* to extract serum. Levels of IFN-γ, CXCL1/KC, TNF, CCL2/MCP-1, IL-12p70, CCL5/RANTES, IL-1β, CXCL10/IP10, GM-CSF, IL-10, IFN-β, IFN-α, IL-6 and IL-4 cytokines were measured using LEGENDplex Mouse Anti-Virus Response (13-plex) Panel (BioLegend) and LEGENDplex Mouse Th (IL-4) Panel (BioLegend) assays. Exported flow cytometry data files were analysed using the LEGENDplex Data Analysis Software (BioLegend).

##### In Vivo Bioluminescence Imaging (IVIS)

Mice were injected intraperitoneally with 150 mg/kg D-luciferin before anaesthesia by isoflurane inhalation (RAS-4 rodent anaesthesia system). After 5 mins, mice were assessed for chemiluminescence using the IVIS Spectrum instrument. For organ imaging, mice were euthanized by cervical dislocation and organs were visualized on the IVIS Spectrum instrument. Mice were exposed for 1 min and organs were exposed for 20 seconds, using field-of-view set to “D”, and with F-stop of 1. Resulting images were analysed using the Living Image software package (Revvity) for enumeration of chemiluminescent signal.

##### CRISPR/Cas9 Knockout of Receptors in OT-I CD8+ T cells

IFNAR1 was knocked out on CD8^+^ OT-I cells following a published method ^89^ using the P3 Primary Cell 4D-Nucleofector®□X Kit S (Lonza). Briefly, 0.3 nmol of each single guide RNA (sgRNA) was mixed with 0.6 µL of Alt-R® S.p.□Cas9□Nuclease V3 (10 mg/mL, Integrated DNA Technologies) and incubated at room temperature for 10 min to form sgRNA/Cas9 ribonucleoprotein (RNP) complexes. 10×10^6^ purified CD8^+^ OT-I cells were resuspended in 10 µL of reconstituted P3 buffer and mixed with sgRNA/Cas9 RNP complex prior to electroporation using a Lonza 4D-Nucleofector (program code: DN100). Electroporated cells were recovered in complete RPMI media (10% FBS) in a 96-well plate for 10 min at 37 °C. The sgRNAs for *Cd19* (5’-AAUGUCUCAGACCAUAUGGG-3’)□and *Ifnar1* (5’-ACAGUUUCGUGUCAGAGCAG-3’, 5’-AAGCAAGGUGUCACUCAUGG-3’) were purchased from Synthego. 50,000 cells were transferred into recipient mice one day prior to vaccination with mRNA-LPX vaccines encoding Ovalbumin.

MHC Class I Tetramer Production (K^b^-RPL6_120–127_) The H2-K^b^ (K^b^) heavy chain (1-275) with a BirA tag for biotinylation was produced in E. coli cells, as well as the human β2 microglobulin (β2m) as per a previously described method.^90^ The K^b^-RPL6_120–127_ complex was then refolded with 30[mg of heavy chain, 10[mg of β2m and 4[mg of the RPL6_120–127_ peptide.^90^ The final protein was purified and biotinylated before tetramerization.

#### Sporozoite Production and Malaria Challenge

##### Sporozoite Production

Naive Swiss mice were inoculated i.p. with infected RBC from an infected syngeneic donor, with parasitemia then confirmed by Giemsa smear and exflagellation quantified 3 days postinfection. Adult *A. stephensi* mosquitoes were then allowed to feed on anaesthetized mice, and 22 days later sporozoites were dissected from mosquito salivary glands and resuspended in cold PBS.

##### Sporozoite Production

Naive Swiss mice were inoculated i.p. with infected RBC from an infected syngeneic donor, with parasitemia then confirmed by Giemsa smear and exflagellation quantified 3 days postinfection. Adult *A. stephensi* mosquitoes were then allowed to feed on anaesthetized mice, and 22 days later sporozoites were dissected from mosquito salivary glands and resuspended in cold PBS.

##### Sporozoite Challenge

Mice were injected i.v. with 200 or 3000 freshly dissected sporozoites. Blood samples were assessed for parasitemia on days 6, 7, 8, 10 and 12 by flow cytometry after staining with5 μg.mL Hoechst 33258 dye (ThermoFisher) for 1□h at 37□°C. An LSRFortessa (BD Biosciences) with a violet laser (405 nm) was used to excite the dye in infected RBC, and percentages of Hoechst-positive cells were compared with those from uninfected controls. Values of >0.1% were considered to indicate positivity for parasites, and mice positive for two consecutive days were euthanized. Those remaining parasitemia-negative on day 12 were considered to be protected.^18^

### 1. QUANTIFICATION AND STATISTICAL ANALYSES

Treatments were assigned to separately caged litter groups. No statistical methods were used to pre-determine sample sizes, but our sample sizes were similar to those reported in previous publications.^13,18,33^ Numerical data from FlowJo were exported to Excel 16.103 (Microsoft) for calculations of cell counts and percentages. Data are shown as mean values□±□S.D., and data analyzed included outliers unless these could be explained by technical error. Data collection and analysis were not performed blind to the conditions of the experiments. Data were assumed to be normal, but this was not formally tested because the sample sizes used were not well powered for normality testing. Statistical tests are indicated in the figure legends, together with sample sizes (n). Individual P values are indicated in the figures. Significant thresholds: *P□<□0.05, **P□<□0.01, ***P□<□0.001, ****P□<□0.0001. All statistical tests and graphs were produced using Prism v.10 (GraphPad Software).

### 2. KEY RESOURCES TABLE

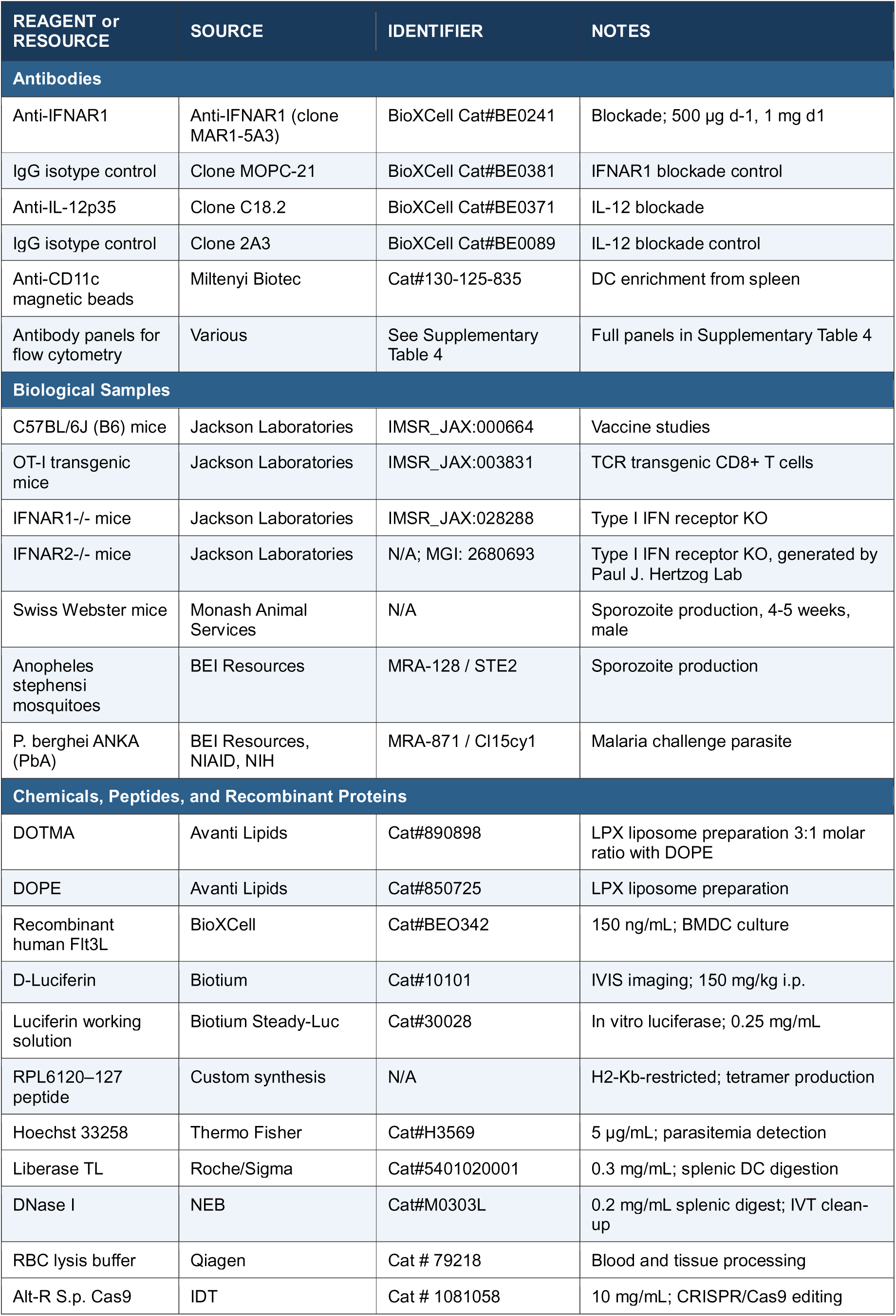
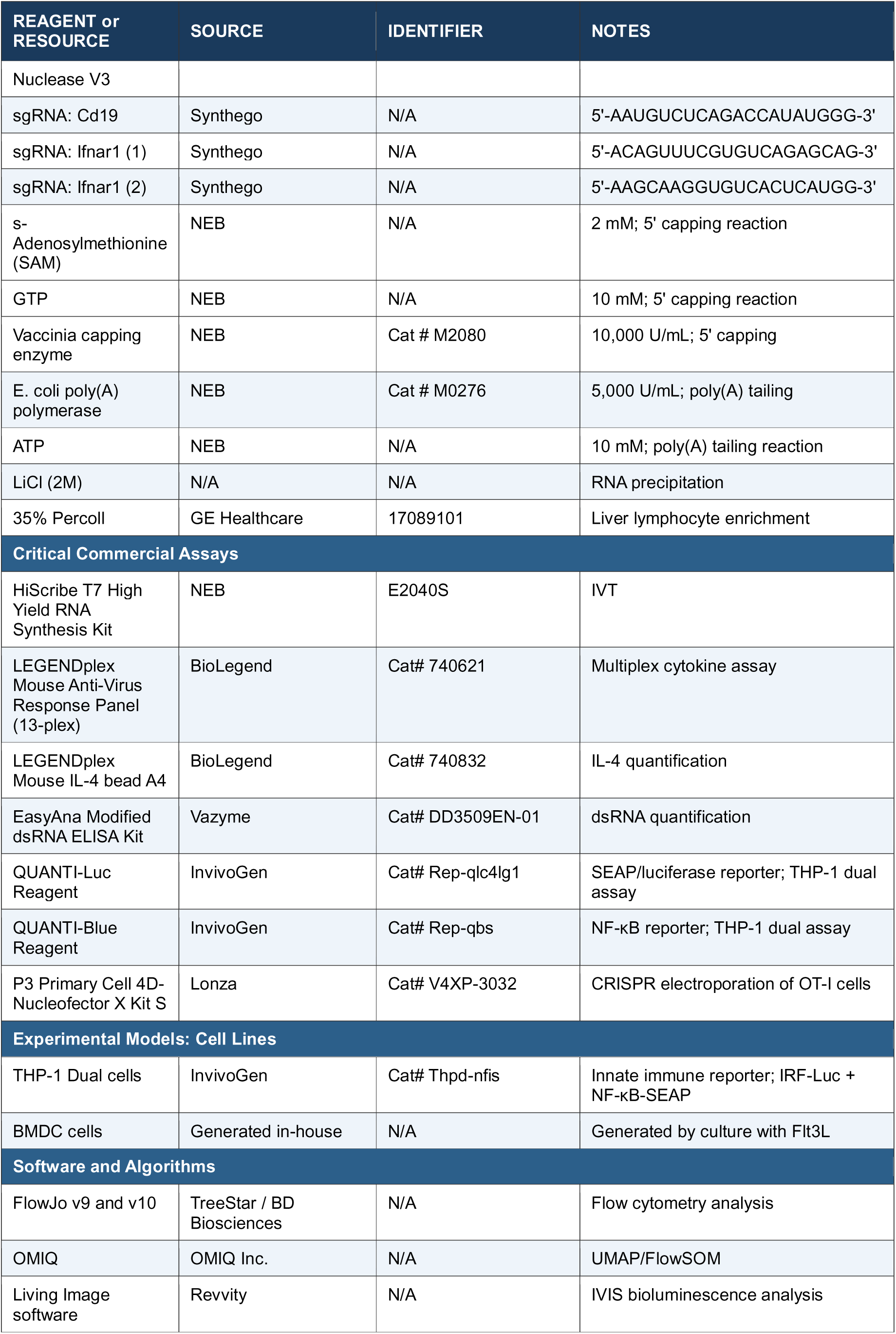
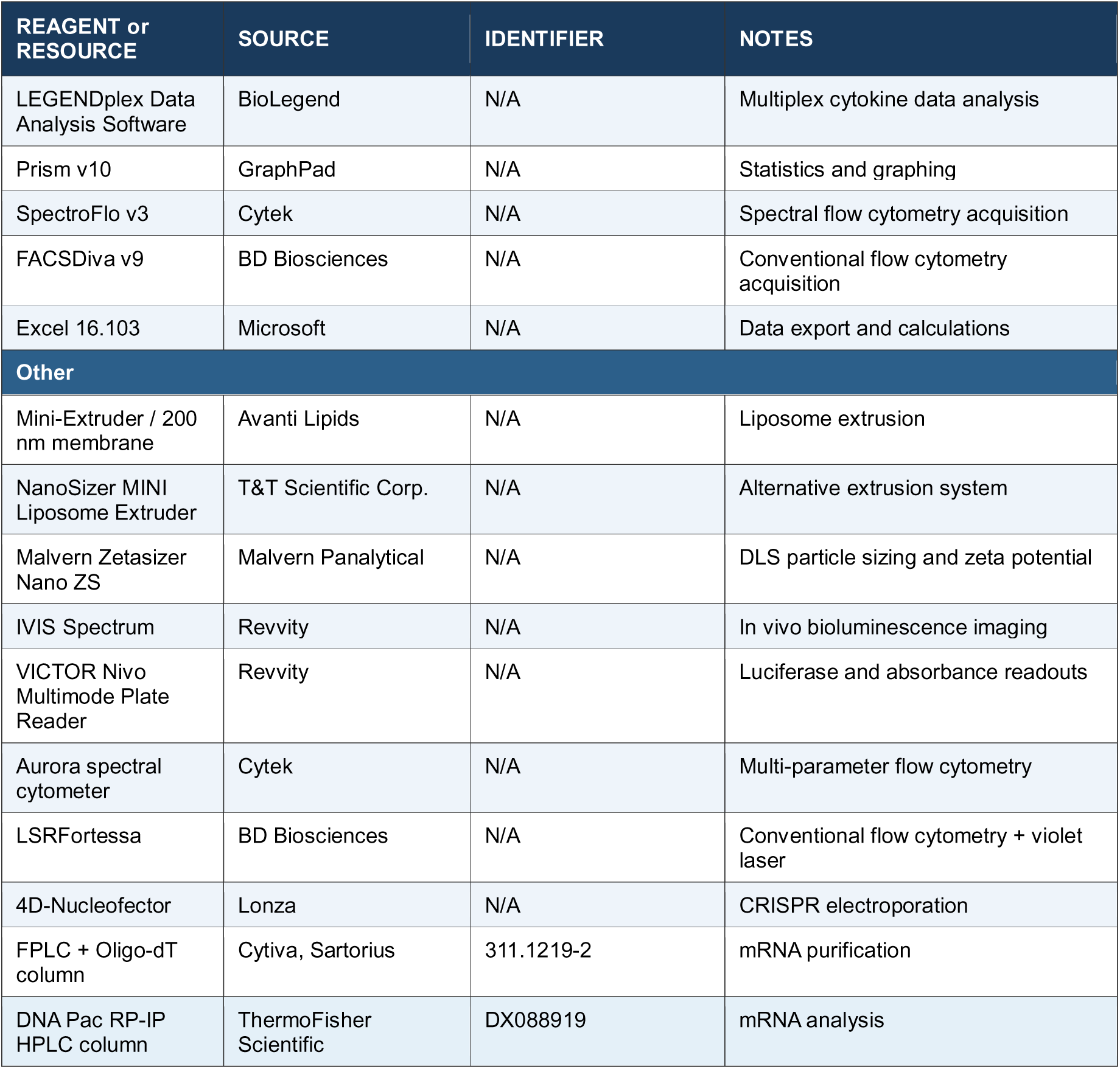

## Data availability

Data to support the findings of study are available from the corresponding author (Prof. Ian Hermans) upon request without restrictions.

## Acknowledgments

We thank the BRF at the Peter Doherty Institute, and the BRU, Hugh Green Cytometry Centre, and NZ RNA platform manufacturing facility at the Malaghan Institute of Medical Research for technical support. We thank D. Godfrey for the CD1d–PBS-44 tetramers and the NIH tetramer core facility for the CD1d-PBS-57 tetramers. J.J.M, K.J.F, O.K.B, J.C.M. and I.F.H were supported by a Ministry of Business Innovation and Employment Independent Research Organisation Fund (MALINSMEDRES2302). J.J.M, E.L, S.L.D, O.R.P, N.C.M, T.B, S.T.S.C, R.J.A, B.J.C, R.E.M, J.N.A.V and G.F.P were supported by a Ministry of Business Innovation and Employment Strategic Scientific Investment Fund (MA-006990).

A.R.L and T.P were supported by the National Health and Medical Research Council of Australia (NHMRC, 2027674). L.H was supported by a Moderna Australia research fellowship and funding from the NHMRC (2027674). S.G and D.J were supported by an NHMRC Leadership Investigator Grant (2034677). A.C and G.I.M were supported by an NHMRC Investigator grant (L3), (GNT2016391). M.G and S.L were supported by the Australian Research Council (ARC DP240102812). L.B was supported by the NHMRC (2038993) and ARC (DP220103545). W.R.H was supported by an NHMRC Investigator fellowship (2033915) and the ARC (DP220103545).

## Author contributions

Conceptualization: J.J.M, M.G Data curation: J.J.M, M.G., A.R.L

Formal analysis: J.J.M, M.G., A.R.L, T.B., E.L, O.K.B. T.P., S.L, S.L.D

Funding acquisition: I.F.H, G.F.P., W.R.H

Investigation: J.J.M, M.G., A.R.L. K.J.F., O.K.B., J.C.M., E.L., S.L., S.L.D., S.T.S.C., T.P

Methodology: J.J.M, M.G., A.R.L, O.R.P., N.C.M., J.C.M., E.L, S.L.D., S.T.S.C., D.J., A.C

Project administration: L.E.H. L.B., I.F.H

Resources: J.N.A.V., R.E.M., I.F.H., G.F.P., W.R.H., G.I.M., S.G, S.L.D., R.J.A., B.J.C

Software: Supervision: I.F.H., G.F.P., L.E.H., W.R.H, L.B., J.J.M

Validation: L.E.H., J.J.M., M.G., A.R.L,

Visualization: J.J.M, A.R.L

Writing – original draft: J.J.M, I.F.H, A.R.L, M.G.

Writing – review and editing: J.J.M, I.F.H., A.R.L, L.E.H., M.G., G.F.P, W.R.H

## Declaration of Generative AI and AI-assisted technologies in the writing process

During the preparation of this work the author(s) used Claude Sonnet V4.6 in order to generate the Abstract, and STAR methods table. After using this tool/service, the author(s) reviewed and edited the content as needed and take(s) full responsibility for the content of the publication.

