## Supplemental Information for "Optimization of a liver Trm cell-inducing mRNA vaccine by reduction of type I interferon response"

**Supplementary Table 1: Analyte concentrations after LPX vaccination for Supplementary Figure 2.** Analyte concentrations were measured in the serum of mice at time-points indicated: Mean ± S.D values are calculated for each time-point. Data were compared by 2-way mixed effects analysis with Sidak's multiple comparisons test. P values are provided for each comparison. Significant P values (<0.05) are bolded.

| *Vaccine Treatment* | | *1h* | *3h* | *6h* | *9h* | *12h* | *24h* |
|---|---|---|---|---|---|---|---|
| ***Data expressed as pg/mL values represented as mean (SD) from two biological repeats*** | | | | | | | |
| *LPX* | IFN$\alpha$ | 44 (47) | 592 (665) | 10458 (6529) | 20579 (13523) | 8308 (4813) | 273 (128) |
| *LPX-$\alpha GC_B$* | | 18 (29) | 533 (571) | 6693 (4644) | 21024 (10681) | 5173 (4559) | 678 (456) |
| Adjusted P Value | | 0.6530 | >0.9999 | 0.6398 | >0.9999 | 0.6286 | 0.1238 |
| *LPX* | IFN$\beta$ | 14 (2) | 14 (2) | 80 (99) | 62 (91) | 27 (41) | 14 (2) |
| *LPX-$\alpha GC_B$* | | 14 (2) | 14 (2) | 64 (71) | 204 (217) | 90 (228) | 19 (12) |
| Adjusted P Value | | >0.9999 | 0.9919 | 0.9991 | 0.3953 | 0.9580 | 0.7061 |
| *LPX* | CXCL10 | 3 (3) | 1077 (1004) | 2785 (3507) | 2384 (2590) | 2793 (3878) | 361 (530) |
| *LPX-$\alpha GC_B$* | | 58 (133) | 5540 (3945) | 7304 (1861) | 5026 (2678) | 8211 (12599) | 7736 (7226) |
| Adjusted P Value | | 0.7779 | **0.0350** | 0.8025 | 0.2067 | 0.7765 | 0.0606 |
| *LPX* | CCL2 | 576 (484) | 1262 (281) | 2636 (1483) | 3438 (2118) | 2760 (1859) | 480 (194) |
| *LPX-$\alpha GC_B$* | | 154 (82) | 16818 (3434) | 21945 (3581) | 12502 (3048) | 14637 (8031) | 5745 (3857) |
| Adjusted P Value | | 0.1291 | **<0.0001** | **<0.0001** | **<0.0001** | **0.0063** | **0.0116** |
| *LPX* | IL-6 | 25 (30) | 86 (91) | 186 (258) | 127 (174) | 11 (7) | 12 (12) |
| *LPX-$\alpha GC_B$* | | 21 (23) | 5844 (2248) | 1830 (1266) | 301 (234) | 356 (362) | 169 (178) |
| Adjusted P Value | | 0.9998 | **0.0001** | **0.0153** | 0.3855 | 0.0857 | 0.1235 |
| *LPX* | CCL5 | 11 (2) | 24 (20) | 166 (85) | 185 (86) | 255 (241) | 56 (35) |
| *LPX-$\alpha GC_B$* | | 11 (3) | 47 (13) | 166 (31) | 239 (75) | 293 (286) | 186 (94) |
| Adjusted P Value | | >0.9999 | 0.0616 | >0.9999 | 0.6365 | 0.9997 | **0.0102** |
| *LPX* | IL-10 | 107 (69) | 91 (78) | 133 (100) | 98 (39) | 105 (54) | 99 (50) |
| *LPX-$\alpha GC_B$* | | 54 (29) | 87 (41) | 80 (23) | 123 (55) | 190 (136) | 253 (109 |
| Adjusted P Value | | 0.2536 | >0.9999 | 0.5775 | 0.8311 | 0.4360 | **0.0090** |
| *LPX* | TNF | 149 (169) | 41 (32) | 30 (18) | 28 (19) | 21 (11) | 16 (13) |
| *LPX-$\alpha GC_B$* | | 92 (75) | 930 (178) | 212 (40) | 97 (27) | 89 (60) | 165 (95) |
| Adjusted P Value | | 0.9237 | **<0.0001** | **<0.0001** | **<0.0001** | **0.0340** | **0.0045** |

| | | | | | | | |
|---|---|---|---|---|---|---|---|
| *LPX* | IL-1β | 8 (5) | 11 (8) | 12 (13) | 13 (9) | 13 (9) | 10 (9) |
| *LPX*-$\alpha GC_B$ | | 9 (8) | 19 (11) | 21 (19) | 17 (7) | 17 (7) | 23 (9) |
| Adjusted P Value | | 0.9962 | 0.4024 | 0.8662 | 0.8566 | 0.9387 | **0.0295** |
| *LPX* | GM-CSF | 5 (5) | 5 (5) | 8 (5) | 7 (6) | 12 (16) | 4 (3) |
| *LPX*-$\alpha GC_B$ | | 32 (33) | 395 (99) | 25 (9) | 19 (8) | 16 (7) | 21 (7) |
| Adjusted P Value | | 0.1723 | **<0.0001** | **0.0023** | **0.0132** | 0.9666 | **0.0001** |
| *LPX* | IFNγ | 2 (1) | 2 (1) | 87 (115) | 169 (137) | 28 (27) | 4 (2) |
| *LPX*-$\alpha GC_B$ | | 26 (29) | 1285 (637) | 8256 (5433) | 96658 (5924) | 24823 (8577) | 22323 (5656) |
| Adjusted P Value | | 0.1590 | 0.0008 | 0.0062 | 0.0040 | <0.0001 | <0.0001 |
| *LPX* | IL-12p70 | 2 (1) | 1 (0.4) | 2 (2) | 2 (1) | 2 (0.8) | 1 (0.4) |
| *LPX*-$\alpha GC_B$ | | 2 (0.6) | 103 (117) | 709 (317) | 276 (177) | 24 (13) | 5 (3) |
| Adjusted P Value | | 0.9952 | 0.1290 | **0.0004** | **0.0054** | **0.0029** | **0.0453** |
| *LPX* | CXCL1 | 486 (718) | 505 (183) | 181 (80) | 136 (115) | 98 (65) | 43 (23) |
| *LPX*-$\alpha GC_B$ | | 47 (52) | 2981 (1426) | 491 (226) | 155 (59) | 207 (203) | 260 (127) |
| Adjusted P Value | | 0.4169 | **0.0022** | **0.0103** | 0.9984 | 0.5896 | **0.0024** |
| *LPX* | IL-4 | 1 (2) | 3 (5) | 4 (4) | 1 (0.9) | 1 (0.6) | 1 (0.8) |
| *LPX*-$\alpha GC_B$ | | 5 (3) | 741 (199) | 24 (11) | 5 (2) | 6 (3) | 5 (2) |
| Adjusted P Value | | 0.2144 | <0.0001 | 0.0014 | 0.0007 | 0.0050 | 0.0038 |

**Supplementary Table 2: Analyte concentrations following LPX vaccination for Figure 5.** Analyte concentrations were measured in the serum of mice at time-points following vaccination indicated, within 24 hours. Mean ± S.D. were calculated and displayed for each time-point. Data were compared by 2-way mixed effects analysis with Sidak's multiple comparisons test. P values are provided for each comparison. Significant P values (P<0.05) are bolded.

| Vaccine Treatment | | 3h | 6h | 9h | 12h | 24h |
|---|---|---|---|---|---|---|
| ***Data expressed as pg/mL values represented as mean (SD) from two biological repeats*** | | | | | | |
| *Prototype* | IFNα | 12308 (6031) | 100542 (17329) | 100421 (28850) | 42607 (8916) | 2375 (1390) |
| *Optimized* | IFNα | 42 (55) | 446 (383) | 1477 (825) | 1801 (693) | 660 (291) |
| Adjusted P Value | IFNα | **0.0006** | **<0.0001** | **<0.0001** | **<0.0001** | **0.017** |
| *Prototype* | IFNβ | 368. (360.38 ) | 1981 (1168.51) | 1547 (740.80) | 340 (210.85) | Not detected |
| *Optimized* | IFNβ | 3 (7) | 2 (3) | 0.3 (0.7) | 0.5 (1) | Not detected |
| Adjusted P Value | IFNβ | **0.052** | **0.002** | **0.0005** | **0.003** | 0.878 |
| *Prototype* | CXCL10 | 13552 (15326) | 18546 (20367) | 36993 (74917) | 21842 (17374) | 5018 (5594) |
| *Optimized* | CXCL10 | 7294 (8879) | 12257 (11955) | 16896 (15496.) | 4419 (4418) | 4469 (3122) |
| Adjusted P Value | CXCL10 | 0.895 | 0.965 | 0.937 | 0.056 | 0.999 |
| *Prototype* | CCL2 | 180 (69) | 710 (168) | 651 (325) | 724 (401) | 279 (73) |
| *Optimized* | CCL2 | 69 (22) | 115 (37) | 154 (74) | 95 (33) | 128 (36) |
| Adjusted P Value | CCL2 | **0.005** | **<0.0001** | **0.004** | **0.003** | **0.0003** |
| *Prototype* | IL-6 | 9592 (6900) | 5254 (3350) | 937 588) | 712 (479) | 140 (112) |
| *Optimized* | IL-6 | 3445 (2038) | 2202 (1062) | 220 (183) | 180 (142) | 104 (95) |
| Adjusted P Value | IL-6 | 0.102 | 0.093 | **0.018** | **0.032** | 0.946 |
| *Prototype* | CCL5 | 180 (69) | 710 (168) | 651 (325) | 724 (401) | 279 73) |
| *Optimized* | CCL5 | 69 (22) | 115 (37) | 154 (74) | 95 (33) | 128 (36) |
| Adjusted P Value | CCL5 | **0.005** | **<0.0001** | **0.004** | **0.003** | **0.0003** |
| *Prototype* | IL-10 | 114 (78) | 139 (74) | 213 (152) | 382 (327) | 283 (255) |
| *Optimized* | IL-10 | 88 (36) | 86 (55) | 119 (78) | 107 (63) | 151 (86) |

| | | | | | | |
|---|---|---|---|---|---|---|
| Adjusted P Value | | 0.891 | 0.364 | 0.428 | 0.127 | 0.531 |
| *Prototype* | TNF | 1453 (791) | 344 (99) | 188 (45) | 219 (114) | 151 (43) |
| *Optimized* | | 997 (290) | 364 (107) | 180 (68) | 151 (103) | 171 (75) |
| Adjusted P Value | | 0.455 | 0.996 | 0.999 | 0.630 | 0.967 |
| *Prototype* | IL-1b | 45 (31) | 51 (26) | 32 (11) | 55 (47) | 21 (12) |
| *Optimized* | | 25 (16) | 40 (22) | 28 (13) | 27 (20) | 16 (12) |
| Adjusted P Value | | 0.399 | 0.844 | 0.960 | 0.433 | 0.901 |
| *Prototype* | GM-CSF | 311 (152) | 113 (44) | 84 (21) | 88 (21) | 41 (28) |
| *Optimized* | | 375 (108) | 90 (27) | 55 (18) | 60 (23) | 49 (38) |
| Adjusted P Value | | 0.821 | 0.608 | **0.030** | 0.284 | 0.988 |
| *Prototype* | IFNγ | 182 (215) | 2738 (872) | 5086 (2894) | 10244 (4027) | 9571 (5671) |
| *Optimized* | | 768 (308) | 3212 (1170) | 6114 (2638) | 16180 (8058) | 17508 (13234) |
| Adjusted P Value | | 0.998 | 0.853 | 0.932 | 0.254 | 0.487 |
| *Prototype* | IL-12p70 | 136 (83) | 414 (182) | 162 (97) | 69 (36) | 20 (12) |
| *Optimized* | | 179 (175) | 977 (381) | 658 (335) | 142 (50) | 23 (15) |
| Adjusted P Value | | 0.967 | **0.005** | **0.005** | **0.009** | 0.992 |
| *Prototype* | CXCL1 | 2693 (1607) | 544 (156) | 155 (39) | 156 (88) | 158 (61) |
| *Optimized* | | 5791 (1605) | 757 (269) | 175 (57) | 112 (43) | 182 (135) |
| Adjusted P Value | | **0.003** | 0.215 | 0.911 | 0.838 | 0.992 |
| *Prototype* | IL-4 | 5039 (3044) | 1170 (2520) | 91 (79) | 71 (47) | 113 (76) |
| *Optimized* | | 4474 (2343) | 720 (814) | 148 (84) | 117 (85) | 83 (52) |
| Adjusted P Value | | 0.996 | 0.989 | 0.550 | 0.617 | 0.873 |

**Supplementary Table 3**: Antibodies used in this study to evaluate CD8+ memory T cells.

| Antigen | Fluorophore | Identifier (Clone, RRID) | Source | Concentration (μg/mL) |
|---|---|---|---|---|
| CD8 | BUV395 | 53-6.7 AB_2732919 | BD | 0.4 |
| mCD1d tet | BV421 | PBS-57 | NIH tet core | 1:600 |
| CD19 | eF450 | eBio1D3 AB_2734905 | eBioscience | 0.2 |
| CD44 | BV510 | IM7, AB_2650923 | Biolegend | 0.3 |
| NK1.1 | BV650 | PK136, AB_2563159 | Biolegend | 0.7 |
| CXCR6 | BV711 | SA05101, AB_2721558 | Biolegend | 2.0 |
| CX3CR1 | BV786 | SA011F11, AB_2565938 | Biolegend | 0.2 |
| TCRβ | FITC | H57-597, AB_313429 | Biolegend | 1.3 |
| CD4 | AF532 | RM4-5, AB_11218891 | eBioscience | 0.3 |
| CD49a | BB700 | Ha31/8, AB_2861198 | BD | 0.4 |
| PD-1 | PerCP-ef710 | RMP1-30, AB_11151142 | eBioscience | 1.7 |
| NVF tetramer | PE | In house | La Trobe | 1:200 |
| CD69 | Pe-Cy5 | H1.2F3, AB_313113 | Biolegend | 0.5 |
| CD101 | Pe-Cy7 | Moushi101, AB_2573378 | eBioscience | 0.2 |
| KLRG1 | APC | 2F1/KLRG1, AB_10641560 | Biolegend | 1 |
| CD64 | AF647 | X54-5/7.1, AB_2566561 | Biolegend | 2.5 |
| CD62L | AF700 | MEL-14, AB_493719 | Biolegend | 0.6 |
| CD45.2 | APC-Fire750 | 104, AB_2629723 | Biolegend | 0.5 |

**Supplementary Table 4:** Antibodies used in this study to evaluate blood CD8 T cells.

| Antigen | Fluorophore | Identifier (Clone, RRID) | Source | Concentration (µg/mL) |
|---|---|---|---|---|
| CD8 | BUV395 | 53-6.7 AB_2732919 | BD | 0.4 |
| CD19 | eF450 | eBio1D3 AB_2734905 | eBioscience | 0.2 |
| CD44 | BV510 | IM7, AB_2650923 | Biolegend | 0.3 |
| CX3CR1 | BV786 | SA011F11, AB_2565938 | Biolegend | 0.2 |
| TCRβ | FITC | H57-597, AB_313429 | Biolegend | 1.3 |
| NVF tetramer | PE | In house | La Trobe | 1:200 |
| CD69 | Pe-Cy5 | H1.2F3, AB_313113 | Biolegend | 0.5 |
| CD127 | Pe-Cy7 | 7R34, AB_1937265 | Biolegend | 0.7 |
| KLRG1 | APC | 2F1/KLRG1, AB_10641560 | Biolegend | 1 |
| CD62L | AF700 | MEL-14, AB_493719 | Biolegend | 0.6 |
| CD45.2 | APC-Fire750 | 104, AB_2629723 | Biolegend | 0.5 |

**Supplementary Table 5:** Antibodies used in this study to evaluate splenic dendritic cells.

| Antigen | Fluorophore | Identifier (Clone, RRID) | Source | Concentration (μg/mL) |
|---|---|---|---|---|
| CD19 | BUV395 | 1D3, AB_2722495 | BD | 0.6 |
| TCRβ | BUV737 | H57-597, AB_2870145 | BD | 0.5 |
| MHC-II (IA/IE) | BUV805 | 269, AB_2873111 | BD | 2 |
| CD8α | BV421 | 53-6.7, AB_2738474 | BD | 0.2 |
| Ly6C | eF450 | HK1.4, AB_10805519 | eBioscience | 0.3 |
| Ly6G | BV570 | 1A8, AB_10899738 | Biolegend | 0.5 |
| BST2 (CD317) | BV650 | 927, AB_2744173 | BD | 1 |
| Sirp1a (CD172α) | BV711 | P84, AB_2740429 | BD | 1 |
| CCR7 (CD197) | BV786 | 4B12, AB_2738765 | BD | 4 |
| CD4 | AF532 | RM4-5, AB_11218891 | eBioscience | 0.2 |
| CD11b | BB700 | M1/70, AB_2744272 | BD | 0.3 |
| XCR1 | PE | ZET, AB_2563843 | Biolegend | 0.4 |
| Ly6A/E (Sca-1) | PE-CF594 | D7, AB_2737751 | BD | 0.1 |
| CD11c | PE-Cy7 | HL3, AB_647251 | BD | 1 |
| CD80 | APC | 16-10A1, AB_1645212 | BD | 0.7 |
| CD86 | APC-R700 | GL1, AB_2739258 | BD | 0.3 |
| CD45.2 | APC-Fire750 | 104, AB_2629723 | Biolegend | 0.5 |

**Supplementary Table 6:** Template DNA sequences used in this study. T7 promotor sequences are underlined.

**Prototype OVA:**

taatacgactcactatagggagacccaagctggctagcgtttaaacttaagcttggtaccgagctcggatccactagtccagtgtggtggaattcgccaccatgggcagcatcggcgccgccagcatggagttctgcttcgacgtgttcaaggagctgaaggtgcaccacgccaacgagaacatcttctactgccccatcgccatcatgagcgccctggccatggtgtacctgggcgccaaggacagcaccaggacccagatcaacaaggtggtgaggttcgacaagctgcccggcttcggcgacagcatcgaggcccagtgcggcaccagcgtgaacgtgcacagcagcctgagggacatcctgaaccagatcaccaagcccaacgacgtgtacagcttcagcctggccagcaggctgtacgccgaggagaggtaccccatcctgcccgagtacctgcagtgcgtgaaggagctgtacaggggcggcctggagcccatcaacttccagaccgccgccgaccaggccagggagctgatcaacagctgggtggagagccagaccaacggcatcatcaggaacgtgctgcagcccagcagcgtggacagccagaccgccatggtgctggtgaacgccatcgtgttcaagggcctgtgggagaaggccttcaaggacgaggacacccaggccatgcccttcagggtgaccgagcaggagagcaagcccgtgcagatgatgtaccagatcggcctgttcagggtggccagcatggccagcgagaagatgaagatcctggagctgcccttcgccagcggcaccatgagcatgctggtgctgctgcccgacgaggtgagcggcctggagcagctggagagcatcatcaacttcgagaagctgaccgagtggaccagcagcaacgtgatggaggagaggaagatcaaggtgtacctgcccaggatgaagatggaggagaagtacaacctgaccagcgtgctgatggccatgggcatcaccgacgtgttcagcagcagcgccaacctgagcggcatcagcagcgccgagagcctgaagatcagccaggccgtgcacgccgcccacgccgagatcaacgaggccggcagggaggtggtgggcagcgccgaggccggcgtggacgccgccagcgtgagcgaggagttcagggccgaccaccccttcctgttctgcatcaagcacatcgccaccaacgccgtgctgttcttcggcaggtgcgtgagcccctga

**Prototype RPL6:**

Taatacgactcactatagggagacccaagctggctagcgtttaaacttaagcttggtaccgagctcggatccgccaccatggccaagaacaccaagagcggcgccagcgccgacgacaagaagaagaccctgaagcactacgtgatcaagggccagaagaagaccctgacccccgtgagggccaagaagaccatcgccaagaagtactacggcaagaagctggccagcaagaagaagtacatcgtgcagaggaagatgaggaagagcatccaggtgggcaaggtggccatcatcctgaccggcaagcacatgggcaagaggtgcatcatcaccaaggtgctgaagagcggcctgctggccgtgatcggcccctacgaggtgaacggcgtgcccctgaagagggtggaccccaggtacctgatcgtgaccagcaccaacgtgttcgacttcaacaacctgagccagatcaaggacaagttcatccaggccgccgagaggatcaacgacgagatcttcatcaagagcatcgacatcaagaagaggcagaagaagctgctgaagaacaagaacgagagcctgttcatgaacgacgtgatccagcagatcaaggagatcagggacagcgaccccaagatgaagaggatcaagctgctgcagaagcagctgggcgacctgctgaagcccgagatcagcaaggacaagatgttcaggagctacatcaagagcaagttcaccctgaggaacaacatgagcttccacaacatcaagttcggcaagcccatccccaacccccctgctgggcctggacagcacctgatga

**Prototype Luciferase**:

Taatacgactcactatagggagacccaagctggctagcgtttaaacttaagcttggtaccgagctcggatccactagtccagtgtggtggaattcgatggaagatgccaagaacatcaagaagggccctgccccattctaccccctggaagatggaacagccggcgagcagctgcacaaggccatgaagagatacgccctggtgcccggcacaatcgccttcaccgatgcccac

atcgaggtggacatcacctacgccgagtacttcgagatgagcgtgcggctggccgaagctatgaagcgctacggcctga
acaccaaccaccggatcgtcgtgtgcagcgagaacagcctgcagttcttcatgcccgtgctgggcgccctgtttatcgg
agtggctgtggcccctgccaacgacatctacaacgagcgcgagctgctgaacagcatgggcatcagccagcccaccg
tggtgttcgtgtccaagaagggactgcagaaaatcctgaacgtgcagaagaagctgcccatcatccagaaaatcatcat
catggacagcaagaccgactaccagggcttccagagcatgtacaccttcgtgaccagccatctgccccctggcttcaa
cgagtacgacttcgtgcccgagagcttcgaccgggacaagacaatcgccctgatcatgaacagcagcggcagcaccg
gactgcctaaaggcgtggccctgcctcacagaactgcctgcgtgcggtttagccacgcccgggaccctatcttcggcaa
ccagatcatccccgacaccgccatcctgagcgtggtgcctttccaccacggcttcggcatgttcaccaccctgggctac
ctgatctgcggcttccgggtggtgctgatgtacagattcgaggaagaactgttcctgcggagcctgcaggactacaagat
ccagagcgccctgctggtgcctaccctgttcagcttctttgccaagagcaccctgatcgataagtacgacctgagcaac
ctgcacgagatcgcctctggcggagccccccctgtctaaagaagtgggagaggccgtggccaagcggttccatctgcctg
gcatcagacagggctatggcctgaccgagacaaccagcgccattctgatcacccccgagggcgacgataagcctggc
gccgtgggaaaggtggtgccattcttcgaggccaaggtggtggacctggacaccggcaagacactgggcgtgaaccag
aggggcgaactgtgtgtgcggggacctatgatcatgagcggctacgtgaacaaccccgaggccaccaacgccctgatt
gacaaggatggctggctgcacagcggcgacattgcctactgggacgaggacgagcacttcttcatcgtggaccggctga
agtccctgatcaagtacaagggctaccaggtggccccagccgagctggaatctatcctgctgcagcaccccaacatct
tcgatgccggcgtggcaggactgcccgatgatgatgctggcgaactgccagccgctgtggtggtgctggaacacggaaa
gaccatgaccgagaaagaaatcgtggactacgtggccagccaagtgaccaccgccaagaaactgagaggcggcgtg
gtgtttgtggacgaggtgccaaagggcctgacaggcaagctggacgcccggaagatcagagagatcctgattaaggcc
aagaaaggcggcaagatcgccgtgtga

**Optimized RPL6**

aatacgactcactataAgccaccatggccaagaacaccaagagcggcgccagcgccgacgacaagaagaagacc
ctgaagcactacgtgatcaagggccagaagaagaccctgacccccgtgagggccaagaagaccatcgccaagaagt
actacggcaagaagctggccagcaagaagaagtacatcgtgcagaggaagatgaggaagagcatccaggtgggcaa
ggtggccatcatcctgaccggcaagcacatgggcaagaggtgcatcatcaccaaggtgctgaagagcggcctgctggc
cgtgatcggcccctacgaggtgaacggcgtgcccctgaagagggtggaccccaggtacctgatcgtgaccagcacca
acgtgttcgacttcaacaacctgagccagatcaaggacaagttcatccaggccgccgagaggatcaacgacgagatct
tcatcaagagcatcgacatcaagaagaggcagaagaagctgctgaagaacaagaacgagagcctgttcatgaacgac
gtgatccagcagatcaaggagatcagggacagcgaccccaagatgaagaggatcaagctgctgcagaagcagctggg
cgacctgctgaagcccgagatcagcaaggacaagatgttcaggagctacatcaagagcaagttcaccctgaggaaca
acatgagcttccacaacatcaagttcggcaagcccatccccaacccccctgctgggcctggacagcacctgatga

**Optimized Luciferase**

TaatacgactcactataAgatggaagatgccaagaacatcaagaagggccctgccccattctacccccctggaagatgg
aacagccggcgagcagctgcacaaggccatgaagagatacgccctggtgcccggcacaatcgccttcaccgatgcc
cacatcgaggtggacatcacctacgccgagtacttcgagatgagcgtgcggctggccgaagctatgaagcgctacggc
ctgaacaccaaccaccggatcgtcgtgtgcagcgagaacagcctgcagttcttcatgcccgtgctgggcgccctgtttat
cggagtggctgtggcccctgccaacgacatctacaacgagcgcgagctgctgaacagcatgggcatcagccagccca

ccgtggtgttcgtgtccaagaagggactgcagaaaatcctgaacgtgcagaagaagctgcccatcatccagaaaatcat
catcatggacagcaagaccgactaccagggcttccagagcatgtacaccttcgtgaccagccatctgccccctggctt
caacgagtacgacttcgtgcccgagagcttcgaccgggacaagacaatcgccctgatcatgaacagcagcggcagca
ccggactgcctaaaggcgtggccctgcctcacagaactgcctgcgtgcggtttagccacgcccgggaccctatcttcgg
caaccagatcatccccgacaccgccatcctgagcgtggtgcctttccaccacggcttcggcatgttcaccaccctggg
ctacctgatctgcggcttccgggtggtgctgatgtacagattcgaggaagaactgttcctgcggagcctgcaggactaca
agatccagagcgccctgctggtgcctaccctgttcagcttctttgccaagagcaccctgatcgataagtacgacctgagc
aacctgcacgagatcgcctctggcggagccccctgtctaaagaagtgggagaggccgtggccaagcggttccatctg
cctggcatcagacagggctatggcctgaccgagacaaccagcgccattctgatcacccccgagggcgacgataagcc
tggcgccgtgggaaaggtggtgccattcttcgaggccaaggtggtggacctggacaccggcaagacactgggcgtgaa
ccagaggggcgaactgtgtgtgcggggacctatgatcatgagcggctacgtgaacaaccccgaggccaccaacgccc
tgattgacaaggatggctggctgcacagcggcgacattgcctactgggacgaggacgagcacttcttcatcgtggaccg
gctgaagtccctgatcaagtacaagggctaccaggtggccccagccgagctggaatctatcctgctgcagcaccccaa
catcttcgatgccggcgtggcaggactgcccgatgatgatgctggcgaactgccagccgctgtggtggtgctggaacacg
gaaagaccatgaccgagaaagaaatcgtggactacgtggccagccaagtgaccaccgccaagaaactgagaggcgg
cgtggtgtttgtggacgaggtgccaaagggcctgacaggcaagctggacgcccggaagatcagagagatcctgattaag
gccaagaaaggcggcaagatcgccgtgtga

Supp. Fig 1

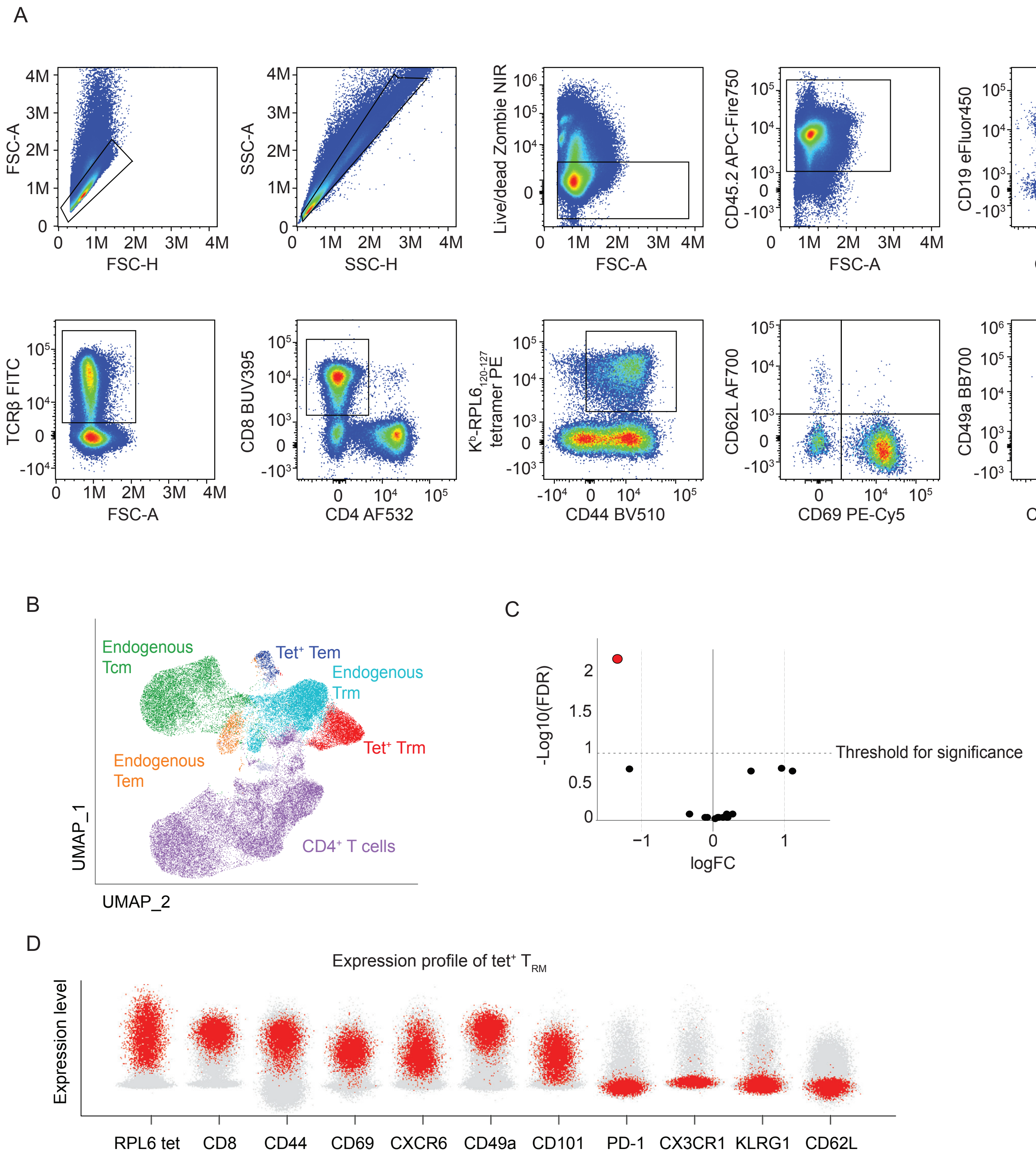

A
FSC-A
FSC-H
SSC-A
SSC-H
Live/dead Zombie NIR
FSC-A
CD45.2 APC-Fire750
FSC-A
CD19 eFluor450
CD64 AF647
TCRβ FITC
FSC-A
CD8 BUV395
CD4 AF532
K^b-RPL6 120-127 tetramer PE
CD44 BV510
CD62L AF700
CD69 PE-Cy5
CD49a BB700
CXCR6 BV711
B
Endogenous Tcm
Tet+ Tem
Endogenous Trm
Tet+ Trm
Endogenous Tem
CD4+ T cells
UMAP_1
UMAP_2
C
-Log10(FDR)
logFC
Threshold for significance
D
Expression profile of tet+ TRM
Expression level
RPL6 tet
CD8
CD44
CD69
CXCR6
CD49a
CD101
PD-1
CX3CR1
KLRG1
CD62L

**Supplementary data 1: High dimensional flow cytometry analysis of liver T cells reveals IFN-I at vaccine priming specifically inhibits liver Trm generation.** (A) Flow cytometry gating strategy used to assess and liver and splenic T cell populations throughout this work, including in panels B-D. (B) Uniform manifold projection (UMAP) showing clusters generated with OMIQ package from flow cytometry analysis of liver T cells 28 day after vaccination with RPL6-LPX-αGC$_B$ in B6 versus IFNAR1ΔCD11c mice (n=10 per group). Clusters are based on cell phenotype as defined by marker expression. Antibody panel using for analysis is provided in Supplementary Table 3, which included H-K$^b$-RPL6$_{120-127}$ tetramers (Tet), and gating strategy to gather T cells is presented in panel A above. (C) Volcano plot comparing cluster size (cell count) between vaccinated WT and IFNAR1ΔCD11c mice. Significance is annotated when the base-10 logarithm of the false discovery rate (log10 FDR) value exceeds indicated threshold, defined by the EDGR algorithm within the OMIQ server. The RPL6-specific Trm cell cluster (Tet+ Trm), which is significantly increased in IFNAR1ΔCD11c mice, is coloured red. (D) Cell surface marker expression profile of Tet+ Trm cluster (red) imposed on marker distribution across all T cells in the dataset (grey).

Supp. Fig 2

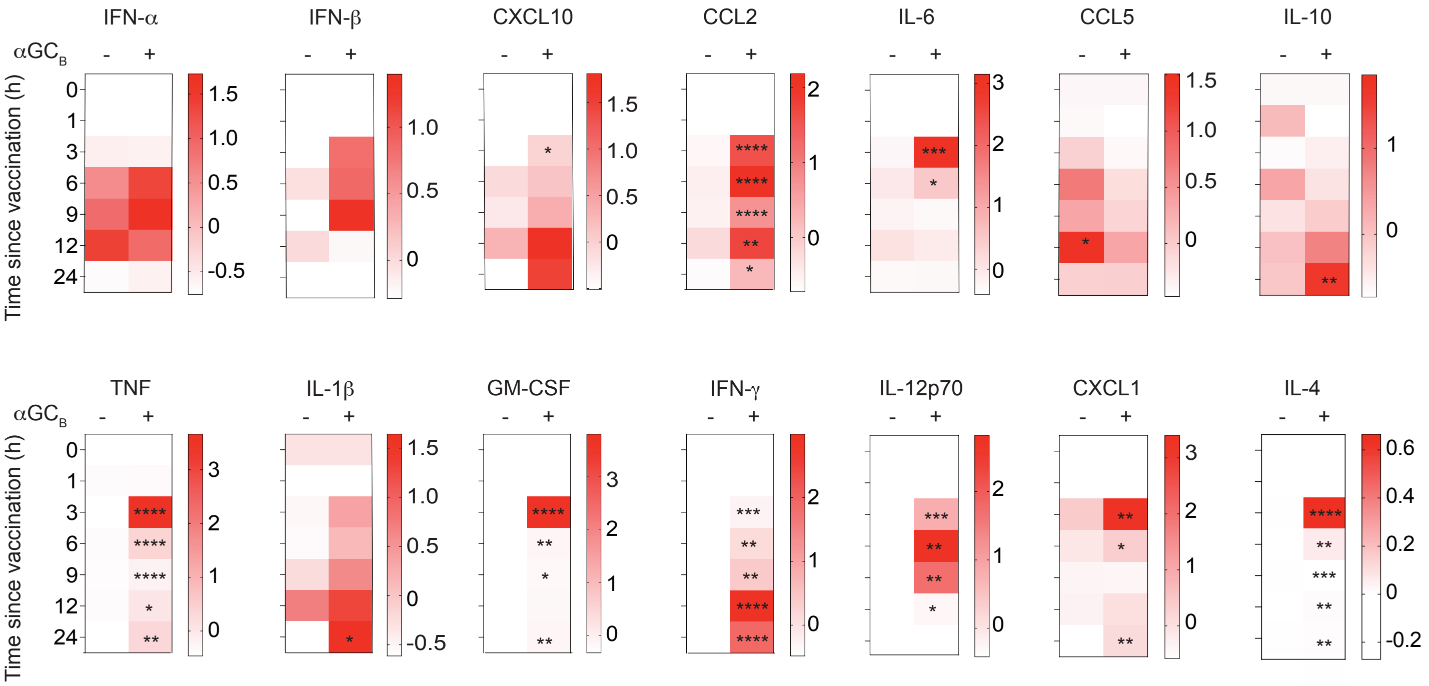
IFN-α
IFN-β
CXCL10
CCL2
IL-6
CCL5
IL-10
TNF
IL-1β
GM-CSF
IFN-γ
IL-12p70
CXCL1
IL-4
αGC$_B$
Time since vaccination (h)

**Supplementary data 2: Inclusion of $\alpha GC_B$ adjuvant enhances serum cytokine and chemokine following vaccination with LPX.** Z-score plots for each analyte representative of relative concentration in the serum at indicated time-points after vaccination with OVA-LPX with or without $\alpha GC_B$. (n=10 per group). Analyte concentrations are provided in Supplementary Table 1. Data were compared by 2-way mixed effects model followed by Sidak's multiple comparisons test. P values are provided in Supplementary table 2 and also indicated on the Z-score plots: (*P < 0.05, **P < 0.01, ***P < 0.001, ****P < 0.0001).

Supp. Fig 3

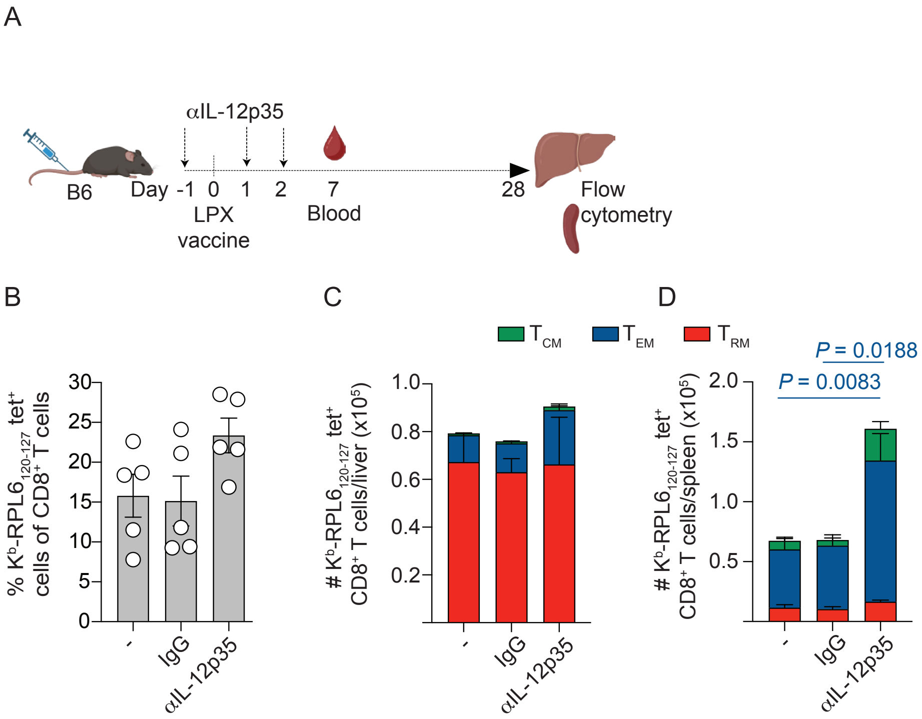
A
αIL-12p35
B6
Day -1 0 1 2 7 28
LPX vaccine
Blood
Flow cytometry
B
% Kb-RPL6120-127 tet+ cells of CD8+ T cells
30
25
20
15
10
5
0
-
IgG
αIL-12p35
C
TCM
TEM
TRM
### Kb-RPL6120-127 tet+ CD8+ T cells/liver (x105)
1.0
0.8
0.6
0.4
0.2
D
P = 0.0188
P = 0.0083
### Kb-RPL6120-127 tet+ CD8+ T cells/spleen (x105)
2.0
1.5
1.0
0.5

**Supplementary data 3: IL-12 is not involved in vaccine-induced liver Trm induction.** IL-12 was blocked using monoclonal antibody against IL-12p35 or isotype control one day prior, and one day after, LPX vaccination with optimised vaccine. (A) Mice were treated with respective antibodies as indicated relative to vaccination (n = 5; derived from one experiment). (B) Kb-RPL6$_{(120-127)}$-specific CD8 T cells were enumerated in the blood, compared by Students T test. K$^{b}$-RPL6$_{(120-127)}$-specific Tcm, Tem and Trm cells in the liver (C) and spleen (D) were enumerated. Data are displayed as mean ± S.D. and were log transformed and compared by one-way ANOVA with Tukey's multiple comparison post-test (P values indicated).

Supp. Fig 4

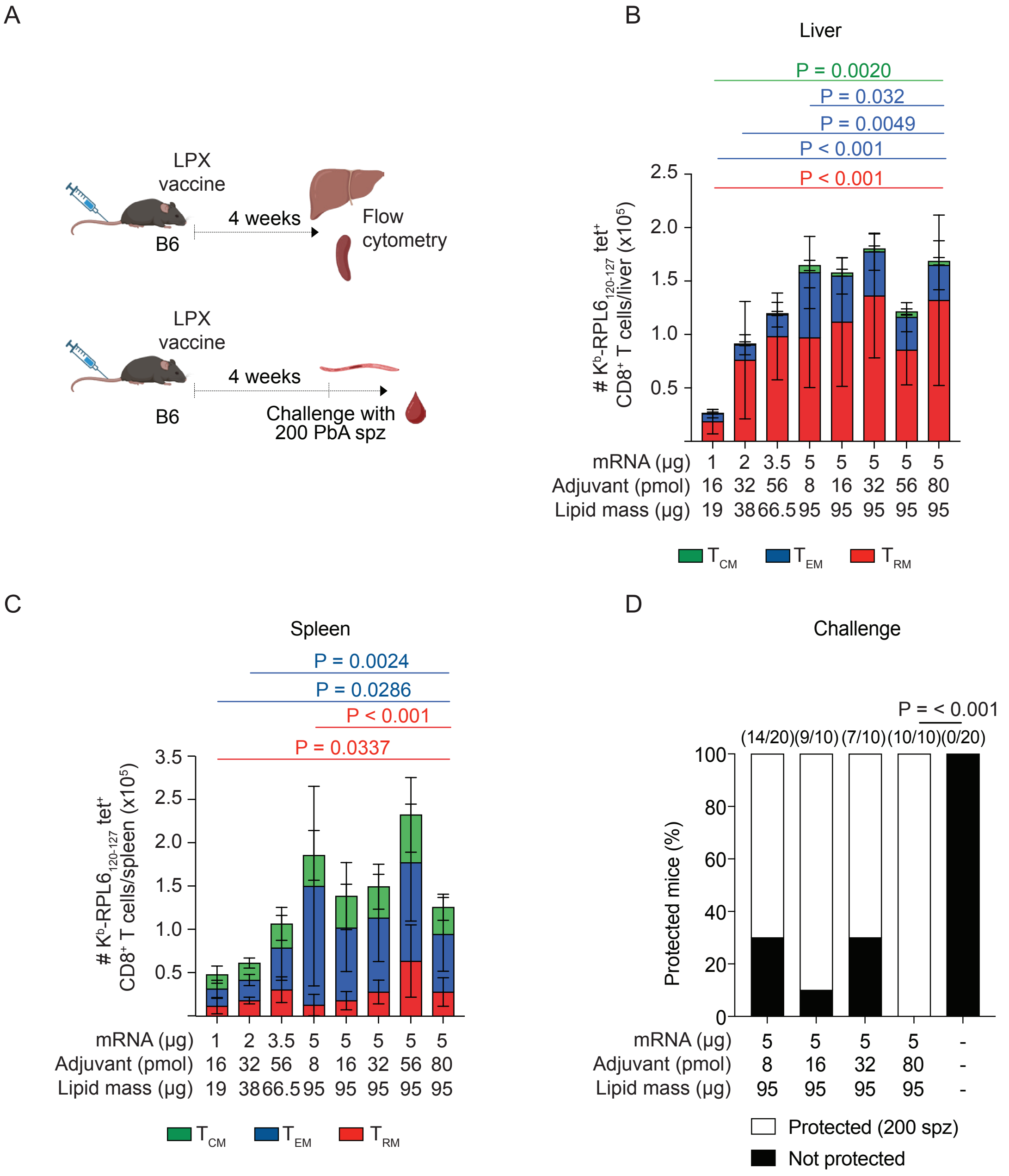
A
LPX vaccine
B6
4 weeks
Flow cytometry
LPX vaccine
B6
4 weeks
Challenge with 200 PbA spz
B
Liver
P = 0.0020
P = 0.032
P = 0.0049
P < 0.001
P < 0.001
### K^b-RPL6 120-127 tet+ CD8+ T cells/liver (x10^5)
mRNA (µg) 1 2 3.5 5 5 5 5 5
Adjuvant (pmol) 16 32 56 8 16 32 56 80
Lipid mass (µg) 19 38 66.5 95 95 95 95 95
TCM TEM TRM
C
Spleen
P = 0.0024
P = 0.0286
P < 0.001
P = 0.0337
### K^b-RPL6 120-127 tet+ CD8+ T cells/spleen (x10^5)
mRNA (µg) 1 2 3.5 5 5 5 5 5
Adjuvant (pmol) 16 32 56 8 16 32 56 80
Lipid mass (µg) 19 38 66.5 95 95 95 95 95
TCM TEM TRM
D
Challenge
P = < 0.001
(14/20) (9/10) (7/10) (10/10) (0/20)
Protected mice (%)
mRNA (µg) 5 5 5 5 -
Adjuvant (pmol) 8 16 32 80 -
Lipid mass (µg) 95 95 95 95 -
Protected (200 spz)
Not protected

**Supplementary data 4: Optimal vaccine dose and adjuvant concentration. Mice were administered one dose of vaccine**. Memory responses were measured in the liver and spleen in a subset of mice 4 weeks later. The remaining mice were challenged with a low dose of 200 sporozoites (spz). (A) Mice were vaccinated with optimised vaccine comprising of the amount of mRNA, $\alpha GC_B$, and lipid as stated. $K^b$-RPL6120-127-specific Tcm, Tem and Trm cells in the liver and spleen were enumerated. (n = 6 – 14; derived from 2 – 6 independent experiments). Data are displayed as mean ± S.D. and were log-transformed and compared by one-way ANOVA with Tukey's multiple comparison post-test (P values indicated). (B) Percentage of mice protected (white) and not protected (black), as measured by parasitemia in the blood up to day 12 post-infection. Numbers above bars indicate proportions of protected mice to the total number of mice. Groups were compared using Fisher's exact test (P values indicated).

Supp. Fig 5

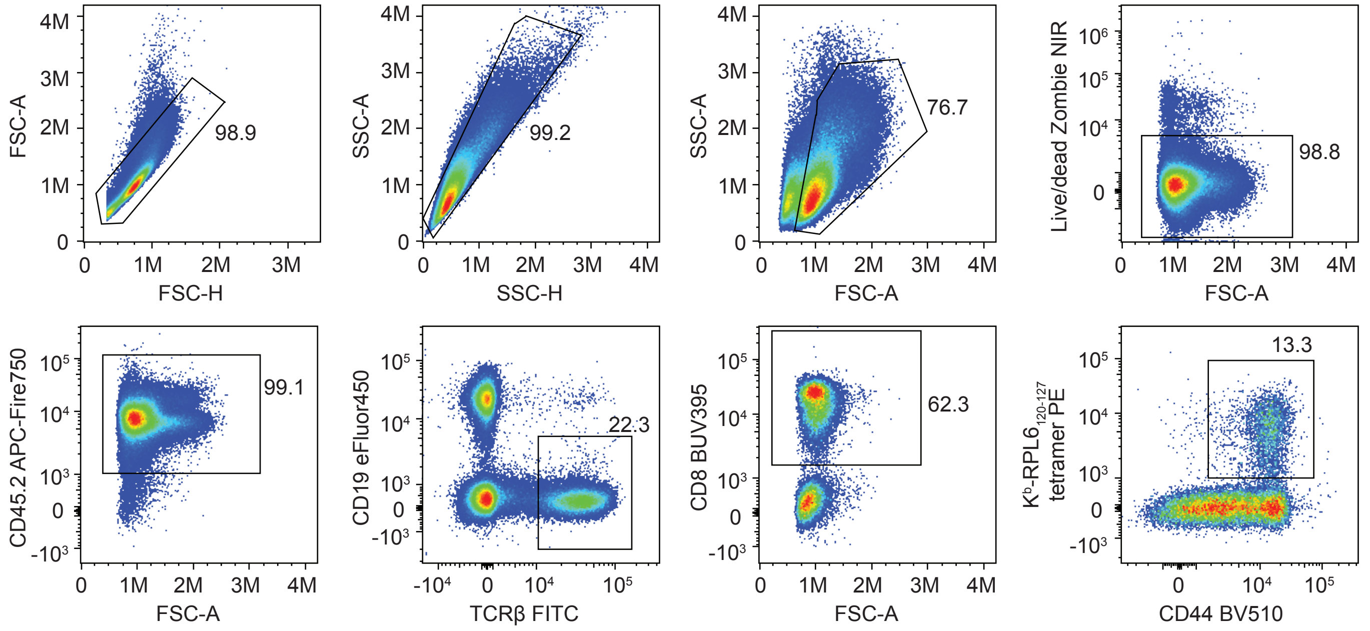
FSC-A
FSC-H
98.9
SSC-A
SSC-H
99.2
SSC-A
FSC-A
76.7
Live/dead Zombie NIR
FSC-A
98.8
CD45.2 APC-Fire750
FSC-A
99.1
CD19 eFluor450
TCRβ FITC
22.3
CD8 BUV395
FSC-A
62.3
K$^b$-RPL6$_{120-127}$ tetramer PE
CD44 BV510
13.3

**Supplementary data 5: Gating strategy used to evaluate blood T cells**. Cells were gated on physical parameters, viability, and surface markers including tetramers

Supp. Fig 6

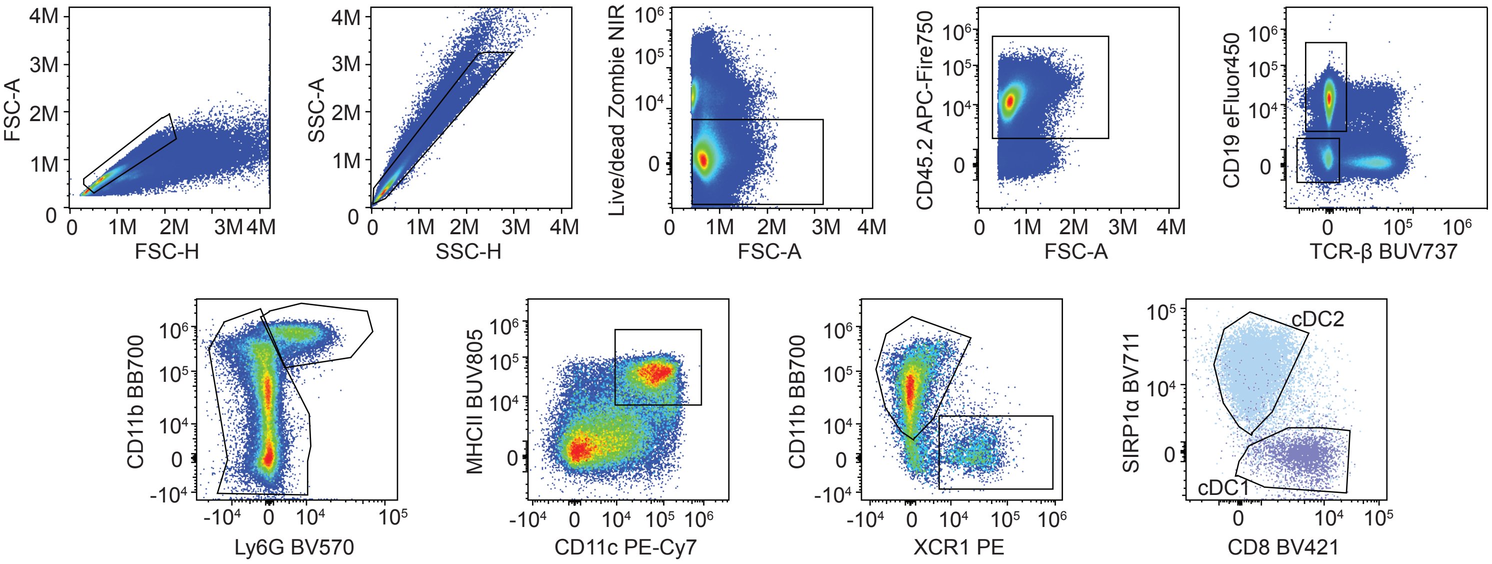
FSC-A
FSC-H
SSC-A
SSC-H
Live/dead Zombie NIR
FSC-A
CD45.2 APC-Fire750
FSC-A
CD19 eFluor450
TCR-β BUV737
CD11b BB700
Ly6G BV570
MHCII BUV805
CD11c PE-Cy7
CD11b BB700
XCR1 PE
SIRP1α BV711
CD8 BV421
cDC2
cDC1

**Supplementary data 6: Gating strategy used to evaluate splenic DCs**. Cells were gated on physical parameters, viability, and surface markers.
